# Body weight, gonadectomy, and other risk factors for diagnosis of osteoarthritis in companion dogs

**DOI:** 10.1101/2023.08.03.550998

**Authors:** Jessica L Graves, Brennen A McKenzie, Zane Koch, Alexander Naka, Nathaniel Spofford, JoAnn Morrison

## Abstract

**OBJECTIVE:** To evaluate age, sex, body weight, breed, neuter status, and age at neutering as risk factors for diagnosis of osteoarthritis in companion dogs

**ANIMALS:** Dogs seen as patients at Banfield Pet Hospital in the United States from 1998-2019 with a date of death in 2019. The final cohort consisted of 131,140 dogs.

**METHODS:** In this retrospective cohort study, Cox proportional hazard models were used to test for associations between osteoarthritis incidence and age at baseline, sex, maximum body weight, maximum body condition score, neuter status, and age at neutering. The same model was used to test these associations in 12 representative breeds, chosen based on breed weight and sample size.

**RESULTS:** Older age, higher adult body weight, gonadectomy, and younger age at gonadectomy were significantly associated with higher risks of osteoarthritis in the total cohort and in all 12 breeds evaluated. Higher body condition score and sex were also significantly associated with osteoarthritis but with minimal effect sizes in the overall cohort, and these risk factors were not consistently significant in all breeds tested.

**CLINICAL RELEVANCE:** These results will assist veterinarians in identifying dogs at higher risk for osteoarthritis and applying appropriate diagnostic, preventative, and treatment interventions. An understanding of potentially modifiable risk factors, such as body condition, and neutering, will support evidence-based discussions with dog owners about risk management in individual patients.

## Introduction

Musculoskeletal pain and lameness are among the most common clinical problems seen in companion dogs.(1–3) A large proportion of these cases involve osteoarthritis (OA), the most common joint disorder in dogs. There is inconsistency in literature regarding the terminology for this condition. Terms such as “osteoarthritis,” osteoarthrosis, and “degenerative joint disease” may be used interchangeably, or some authors may use different terms for conditions with specific etiology or pathogenesis. In this report, we use the term “osteoarthritis” to refer to a diagnosis of one of the following structured diagnostic codes: osteoarthritis, arthritis or degenerative joint disease – excluding rheumatoid, septic or immune mediated arthritic conditions.

Prevalence of OA ranges from 2.5% to over 80% depending on study methods and population characteristics.(1,4,5) This variation likely reflects both true differences in prevalence between populations, related to differences in breed, age, husbandry, and other causal factors, as well as differences in diagnostic methods and case definitions between studies.(6)

As a progressive, incurable condition, OA can affect comfort and quality of life for a substantial proportion of a patient’s life.(4) This condition compromises the welfare of affected dogs and places a significant burden on human caregivers.(7) Musculoskeletal pain and locomotor dysfunction, often due to OA, are also among the most frequent reasons for euthanasia in dogs.(8–11)

Currently, the primary approach to mitigating the negative impact of canine OA is treatment once clinical signs have manifested. Earlier detection and interventions to prevent or delay the development of OA could have significant benefits. Effective approaches for preventing or delaying the development of OA depend on a clear understanding of relevant risk factors and identifying individuals at increased risk.

Many putative risk factors have been associated with development and progression of canine OA.(6) There is strong evidence for the role of genetic factors influencing both individual risk and breed differences in susceptibility to OA. These factors are often associated indirectly with OA, causing predisposition conditions such as hip or elbow dysplasia or a propensity for cranial cruciate ligament rupture, which then lead to development of arthritis.(6,12)

Body weight is another factor associated with OA risk. However, studies often do not clearly distinguish between body size and body condition. Larger breeds appear to be at greater risk, as do individuals who are overweight or obese, but the relationship between these different body-size variables is not always clear.(6,13–15)

Osteoarthritis is considered a disease of aging, and increased age is often associated with increased OA prevalence. This association is potentially complicated, however, by the lack of surveillance and diagnostic markers of early, pre-clinical joint disease. Predisposing conditions and early OA may be present undetected in young dogs while older individuals may be more likely to be diagnosed with OA because of greater diagnostic attention or because the condition has progressed to more apparent clinical signs.(6)

The evidence is limited and conflicting for many potential OA risk actors. Sex, for example, is often associated with OA prevalence, but both males and females have been reported to be at increased risk and the potential for confounding by body size, activity, and neuter status is high.(6)

One of the most debated risk factors for OA is neuter status.(6,16–19) While most reports indicate neutered dogs are at higher risk than intact dogs, the details of the relationship between neuter status and OA are unclear. For example, this association seems to be consistently true for large-breed dogs and less often found in smaller breeds.(18) Neutered dogs are also at greater risk for obesity, so the degree to which the relationship between neutering and OA is mediated or confounded by body condition is uncertain.(6,20–22)

Some studies find that age at neutering influences the impact of gonadectomy on the risk of OA and predisposing conditions, such as cruciate ligament disease.(19,23) This suggests gonadal hormones are protective primarily through effects of skeletal development, and neutering after puberty or skeletal maturity may be less likely to promote OA. However, other studies report residual increased risk in dogs neutered after skeletal maturity and suggest that gonadal hormones may have an ongoing protective effect.(24) There is also significant variation in the existence and strength of associations between neutering and orthopedic disease found in different breeds and different research studies.(18) Many other factors, including diet, activity patterns, and even birth month have been associated with the risk of OA, but detailed causal links have not been clearly identified.(6)

From a preventative-medicine perspective, risk factors for OA can be considered modifiable or non-modifiable. Most genetic factors are not directly modifiable in individuals, and the risk presented by specific genotypes, breed, and conformation are fixed at birth or during development. Recent advances in the study of epigenetics suggest it may be possible to mitigate the impact of some genes through environmental modification of the regulation of gene activity, but clinical interventions for doing so have not yet been validated.(25,26) On a population level, genetic factors which influence the occurrence of OA can potentially be modified by selective breeding targeting both individual genes associated with increased OA risk as well as conformations that predispose to the disease.(25,26)

Other factors associated with OA are clearly modifiable in individuals, including body weight and body condition, diet, activity patterns, and neutering practices.

Common dietary recommendations to delay or prevent OA include reduced feeding to prevent obesity and modulate skeletal development in growing puppies. Lifelong caloric restriction in a cohort of Labrador retrievers delayed the onset of hip OA and reduced the severity of the disorder.(27,28) Less definitive effects were reported for elbow and shoulder OA.(29,30) Whether these effects were due solely to differences in body condition or other influences of caloric restriction is uncertain.

Experimental studies have also shown that reduction in caloric and calcium content of diets can reduce the risk of developmental abnormalities in giant-breed dogs, such as hip dysplasia, that frequently lead to OA later in life.(31,32) Other nutritional interventions may also influence development of OA and predisposing conditions,(33) but there is still significant uncertainty about the efficacy of most dietary approaches.

Greater clarity about the role of key risk factors in development of canine OA would be useful for informing preventative strategies. The purpose of this study was to examine selected risk factors for development of OA in a large, retrospective cohort study of companion dogs using medical record data from primary care veterinary practices. We examined previously reported risk factors including age, sex, breed, body weight, and body condition. We also sought to further investigate the relationships between OA, neuter status and age at neutering as well as the variability of these relationships among breeds.

## Materials and Methods

### Data summary and processing

By the end of 2019 the Banfield Pet Hospital network included 1,084 primary care small animal hospitals in 42 U.S. states, the District of Columbia and Puerto Rico. All hospitals used the same proprietary practice information management system (PIMS; PetWare®) to enter patient and visit information and the resulting electronic medical records were uploaded nightly to a central data warehouse. The PIMS contained both structured and unstructured fields for data entry. Structured data included diagnoses and clinical signs, physical examination findings, and invoiced services and medications; unstructured data included narrative text related to subjective and objective observations, patient assessment and treatment plan.

All in-network visits from dogs with a death date in 2019 were extracted from the data warehouse, with visit information ranging from 1998-2019. Patient information for each visit included breed, sex, neuter status, visit age, and visit weight. Visit information included the location and reason for the visit, all structured clinical signs and diagnoses, examination findings related to musculoskeletal issues and body condition, and any invoiced items or services related to the diagnosis or treatment of osteoarthritis. Invoiced items or services related to in-hospital euthanasia were also extracted in order to more accurately identify patient death date.

To improve data quality the visit-level data was cleaned and evaluated for data entry errors. First, only visits in which the dog received a physical examination by a veterinarian were included. Some data were removed due to likely data entry errors. These include visits in which there were discrepant entries, visit dates occurring after 2019 or weights recorded over 300 lbs, or dogs who had implausible data values (e.g., negative age). To minimize cohort effects, only dogs born after 1997 were included for analysis. Lastly, the body condition score (BCS) assessment has changed over time (e.g., the use of 3-point (clinician-based assessment), 5-point and 9-point BCS scales), scores were harmonized and converted into a single 9-point scale (**Table 1**).

**Table 1:** Body condition score conversion methodology.

| Point scale | Original BCS value | New BCS value | Classified as |
| --- | --- | --- | --- |
| 5-point & 9-point | Body Condition Score - Thin 1 | 1 | Too thin |
| 5-point | Body Condition Score - Underweight 2 | 2 | Too thin |
| 3-point | Nutrition/Weight Control Advice - Thin 1 | 2 | Too thin |
| 9-point | Body Condition Score - Too Thin 2 | 2 | Too thin |
| 9-point | Body Condition Score - Too Thin 3 | 3 | Too thin |
| 9-point | Body Condition Score - Ideal 4 | 4 | Ideal |
| 3-point | Nutrition/Weight Control Advice - Normal 2 | 4 | Ideal |
| 5-point | Body Condition Score - Ideal 3 | 5 | Ideal |
| 9-point | Body Condition Score - Ideal 5 | 5 | Ideal |
| 9-point | Body Condition Score - Above Ideal 6 | 6 | Overweight |
| 5-point | Body Condition Score - Overweight 4 | 7 | Overweight |
| 9-point | Body Condition Score - Overweight 7 | 7 | Overweight |
| 3-point | Nutrition/Weight Control Advice - Heavy 3 | 7 | Overweight |
| 5-point | Body Condition Score - Obese 5 | 8 | Obese |
| 9-point | Body Condition Score - Obese 8 | 8 | Obese |
| 9-point | Body Condition Score - Obese 9 | 9 | Obese |

Osteoarthritis was defined as the earliest diagnosis of the following structured diagnostic codes: osteoarthritis, arthritis or degenerative joint disease – excluding rheumatoid, septic or immune mediated arthritic conditions. Follow-up time was calculated as the number of years between a dog’s age at their first visit and their age at OA diagnosis or death, whichever occurred first – dogs with follow-up times greater than 20 years were removed. In order to capture incidence of aging-related OA, dogs who were not observed into maturity (2 years and up) or who already had OA at their first visit were not included for analysis.

Desexing status was defined based on veterinarian assessment and if the dog was gonadectomized at a Banfield clinic. Age at desexing was defined as age during a visit with a desexing procedure. Age at desexing was only ascertained for dogs desexed at a Banfield clinic. To preserve temporality between desexing as a risk factor and OA incidence, dogs desexed after being diagnosed with OA were treated as intact. Dogs desexed after 5 years of age were not included in age at desexing analyses, as desexing later has a higher likelihood of being a therapeutic rather than an elective procedure.

Multiple weight and BCS metrics were created. To estimate the effects of maximum body size or condition, the 75th-percentile of adult (2+ years of age) body weights and BCS were used to approximate a ‘maximum’ weight or BCS. 75th percentiles were used to minimize the influence of outliers. Median weight and BCS between ages 1.5 and 2.5 years of age were used to approximate the weight and BCS at developmental maturity. Not all dogs were seen between 1.5 and 2.5 years of age. In these cases, weight or BCS at desexing (so long as desexing occurred after full development, e.g., 2+ years of age) was used.

To estimate effects of weight gain after desexing, weight and BCS after desexing was defined as the 75th-percentile of weight and BCS after desexing and 2+ years of age. Percent change in weight and BCS after desexing was calculated as 100*(observation after desexing - observation at maturity)/observation at maturity.

The majority of dogs seen at Banfield are on a wellness plan (a set of pre-paid services meant to enhance the provision of preventive care services). Wellness plan data is structured within the system so the wellness plan variable was converted to a binary variable encoding if a dog ever was on a wellness plan. Dogs whose wellness plan started at or after OA diagnosis were treated as “never” having been on a plan.

### Statistical methods

#### Descriptive statistics

Mean, standard deviation (SD), median, 25th and 75th percentiles, and ranges were calculated for continuous measures: age at baseline, mature and maximum weight, age at OA or censorship/death and duration of follow up (years). Observation counts and proportions were calculated for categorical variables: sex (Female or Male), mature and maximum BCS (1-3=Too thin; 4-5=Ideal; 6-7=Overweight, 8-9=Obese), gonadectomy status (desexed or intact), age at desexing (<6 months, 6-12 months, 12-24 months, 24-60 months), and ever had a Banfield wellness plan (Yes or No). For all variables, the number of missing values were reported. All descriptive statistics are reported for the total cohort as well as stratified by OA diagnosis.

#### Cox proportional hazards

Survival analysis was used to characterize the incidence of OA in companion dogs. The analyses were conducted using duration of follow-up as the measure of survival time, calculated as the years between a dog’s first visit and their OA onset or death. This choice of follow-up time as the survival time scale enables easier estimation of age as a risk factor for OA onset. Dogs were right-censored at death.

A Cox proportional hazards model(34) was used to test joint associations of OA risk factors. In particular, age at baseline, max weight, max BCS, desexing status and sex were used as risk-factors. The Cox model was stratified, separating dogs on (76.3%) and not on (23.7%) wellness plans (**Table 2**). As the dogs on wellness plans were more closely observed on average, the likelihood of OA diagnosis in these dogs was higher (**Supplemental Figure 1**), mostly likely due potential detection bias, and for this reason a stratified Cox proportional hazards model was used. This stratified model accounts for this wellness plan-associated surveillance bias by allowing the baseline hazard of dogs in these groups to be different.(35) On average, dogs on wellness plans visited Banfield 25.06 times (SD 18.54) while dogs not on a wellness plan visited only 4.67 times (SD 6.85).

**Table 2.**
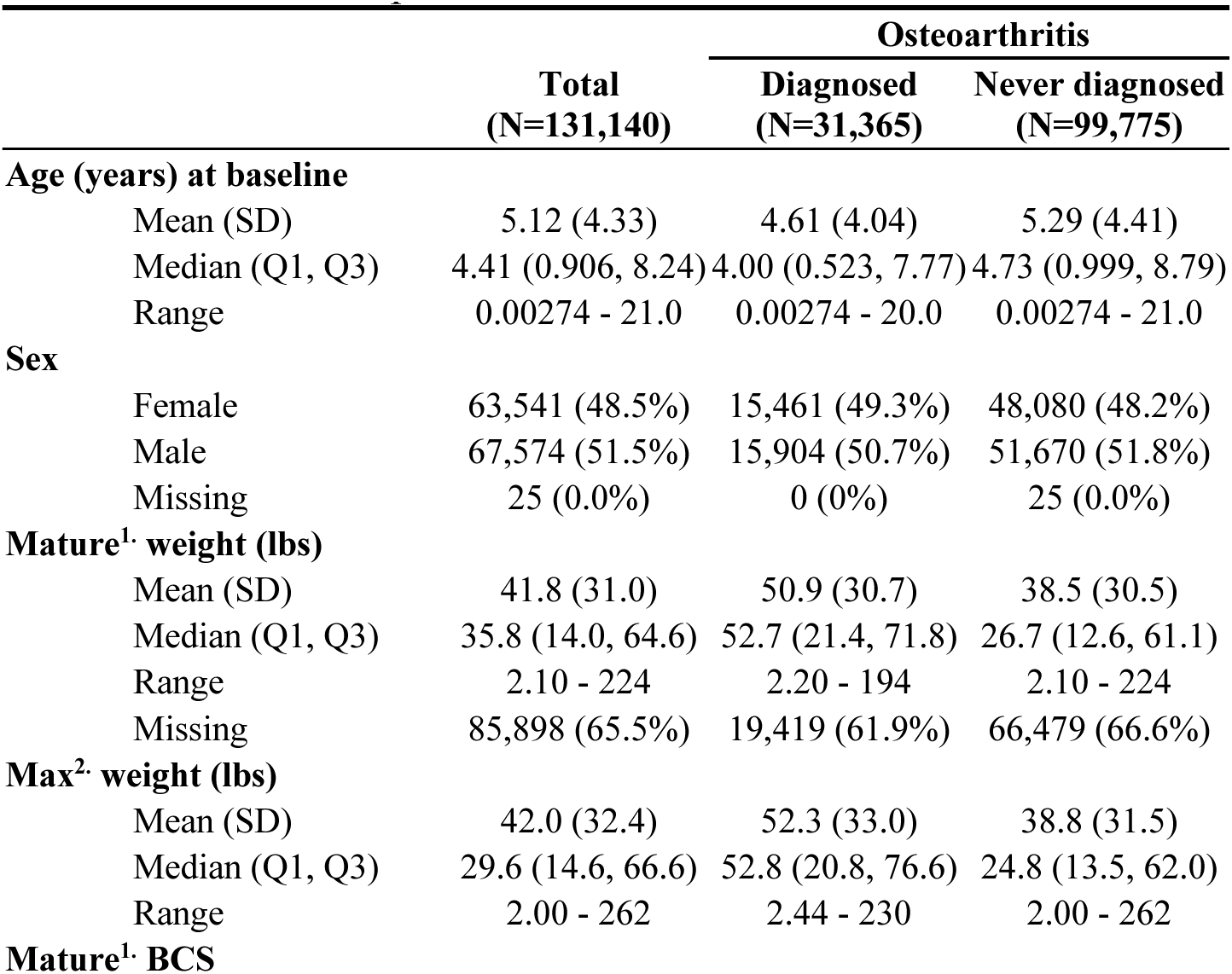
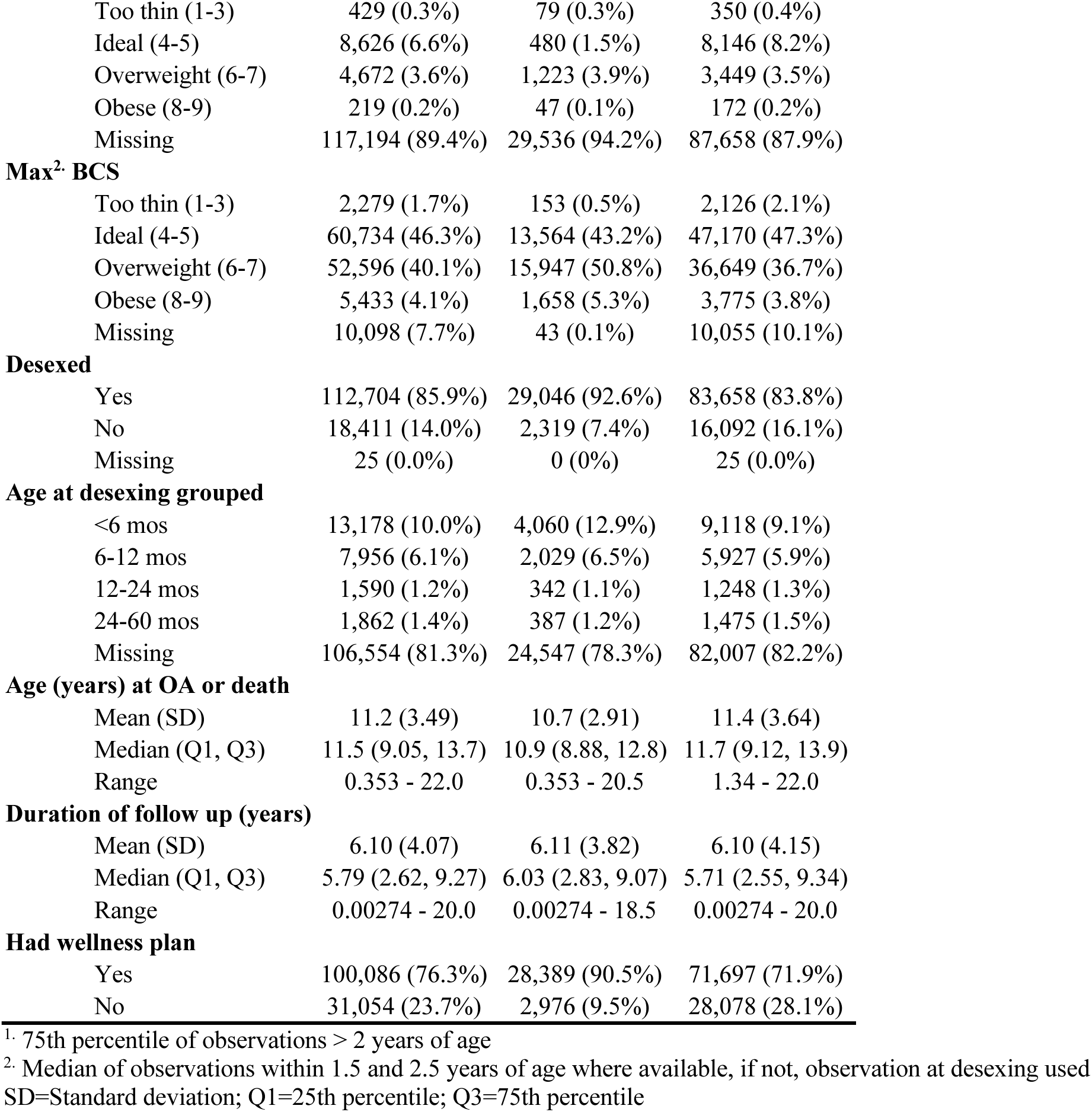
Cohort descriptive statistics.

After fitting the Cox proportional hazards model, using the *coxph* function from the *survival* package(36) in R, the proportional hazards assumption was assessed using *cox.zph* and visualization of residuals. The assumption being met, hazard ratios (HR) and their 95% confidence intervals (CI) were estimated to test the magnitude and direction of each risk-factor on time to OA diagnosis. To account for the risk-factors having varying scales of values, standardized HRs were calculated for continuous risk factors (e.g., age at baseline) by standardizing the risk factor (mean centering and dividing by the standard deviation). For all models, both the natural scale and standardized HRs and 95% CIs are reported, along with concordance index and number of observations.

The marginal effects of each risk-factor on OA-free survival time were plotted to better understand the magnitude of effect sizes. Marginal effects reflect the average effect of a given risk factor, holding all other covariates in the model constant. To visualize effects of continuous risk factors, predicted OA-free survival curves were estimated based on quartile values or clinical relevance. To visualize marginal effects of categorical variables, each level was used to predict OA-free survival curves. For risk factors not being directly visualized, continuous values were assigned as cohort median values (e.g., median weight) and categorical variables were assigned as a proportion (e.g., proportion male). As the majority of dogs were on wellness plans, strata were assigned to being on a wellness plan. For all interaction models, median survival times and their 95% CIs are also reported.

### Breed-specific analyses

For breeds with at least 500 observations, binomial tests of proportion were used to compare breed-specific OA incidence rates against the cohort-wide incidence rate, using Bonferroni adjusted p-value to adjust for multiple comparisons. Given the diversity and size of the cohort, risk factors of OA were also evaluated in 12 representative breeds, chosen based on breed weight and sample size. First, the maximum body weight of all dogs in the cohort was split into quartiles and then breeds were assigned to a weight quartile based on the mean maximum weight of all dogs of that breed. Then, the top three breeds within each quartile with the most samples were chosen for further analysis. This ensures representation of a wide range of dog weights while not including breeds lacking sufficient sample size to make substantive conclusions. The same stratified Cox regression model used to test primary risk factors of OA was applied to each of these 12 breeds.

All analyses were performed in R version 4.1.2. Type I error was set to α = 0.05 for all statistical tests.

## Results

### Descriptive statistics

Descriptive statistics of the final cohort resulting from the data processing steps outlined are reported in **Table 2**. The final cohort included 131,140 dogs, 31,365 (23.9%) of which were diagnosed with OA during the course of the study. The cohort of dogs had an average age entering the study of 5.12 years (SD = 4.33), average mature weight of 41.8 lbs (SD=31.0), and 63,541 (48.5%) dogs were female. There were 318 unique breeds present in the study, with Labrador Retrievers (n=11,718), Chihuahuas (n=10,147) and Yorkshire Terriers (n=6,570) as the most frequent breeds observed.

Not all variables generated during the data processing steps outlined above contain observable data. The relatively high proportion of missing data in mature BCS are due to strict construct definitions to generate this variable. Mature BCS was defined as the median BCS score between 1.5 to 2.5 years – many dogs did not have visits during this window in which BCS was assessed. Similarly, age at desexing could only be quantified for dogs who were gonadectomized at a Banfield hospital, which could be confidently matched to a visit date. Approximately 20% of dogs in this cohort were desexed at Banfield, of which 1% were desexed after receiving an OA diagnosis and 9.4% were desexed > 5 years of age. To preserve temporality and capture patterns of voluntary desexing, age at desexing > 5 years of age or after OA were set to missing and are therefore not included in age at desexing analyses.

In order to fully leverage all available data, only observed data of relevant variables were included in statistical analyses.

### Primary risk factors

To explore if effects of previously reported risk factors were recapitulated in this expansive cohort, we fit stratified Cox proportional hazards model using age at baseline, desexed status (desexed/intact), sex (male/female), adult weight and adult BCS as our primary risk factors, and treated wellness plan as a stratifying variable. Stratification along the presence of being on a wellness plan was supported by evidence of a surveillance (or detection) bias in dogs who were on wellness plans **(Supplemental Figure 1)**.

**Table 3** and **Figure 1** show that older age (Std. HR=3.70 (3.64-3.76)), higher adult body weight (Std. HR=1.83 (1.81-1.85)), and desexing (Std. HR=1.37 (1.31-1.43)) are statistically significantly associated with higher risks of OA in this cohort. While hazard ratios for adult BCS and sex were both statistically significant, these effect sizes are minimal (**Figure 1**).

**Figure 1.**
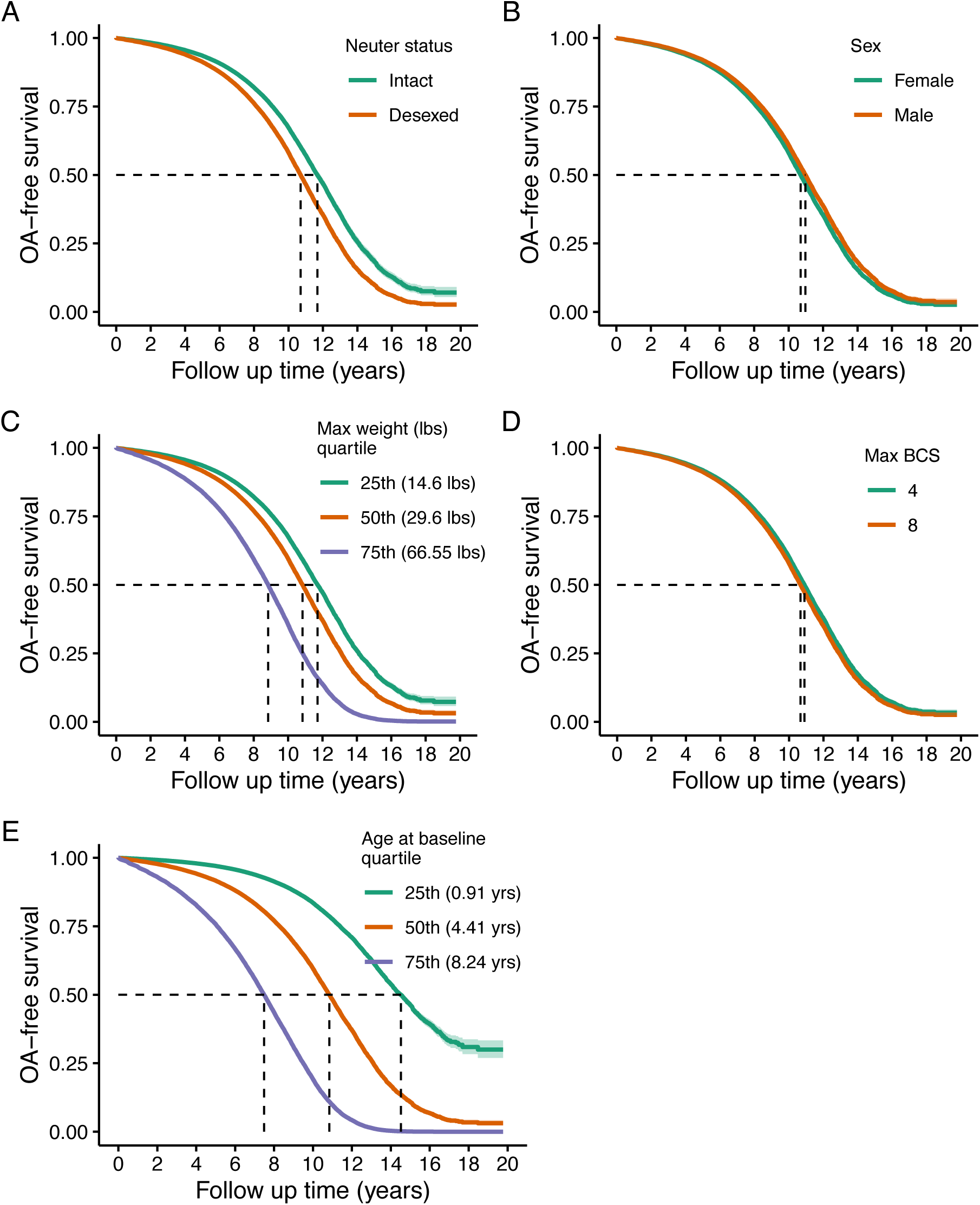
Predicted marginal OA-free survival curves of primary risk factors of OA. Assignments for generating predicted survival curves: median values for continuous covariates not being directly tested, proportions for binary variables, and assuming presence of a wellness plan. Quantiles were used for continuous variable groupings being visualized, except for BCS, which were determined based on clinical relevance.

**Table 3.** Stratified Cox proportional hazard model testing the associations of primary risk factors on OA-free survival.

| Variable | HR (95% CI) | Std. HR (95% CI) | p-value | Concordance | N obs |
| --- | --- | --- | --- | --- | --- |
| Age at baseline | 1.35 (1.35-1.36) | 3.70 (3.64-3.76) | <0.001*** | 0.79 | 121,035 |
| Max weight | 1.02 (1.02-1.02) | 1.83 (1.81-1.85) | <0.001*** |  |  |
| Max BCS | 1.02 (1.01-1.03) | 1.02 (1.01-1.03) | <0.001*** |  |  |
| Desexed | 1.37 (1.31-1.43) | 1.37 (1.31-1.43) | <0.001*** |  |  |
| Male | 0.91 (0.89-0.93) | 0.91 (0.89-0.93) | <0.001*** |  |  |

\*Std. HR= Standardized continuous variables (mean centered, scaled by standard deviation). Wellness plan treated as stratifying variable.

### Breed-specific risk factors: Are risk factors consistent within breeds?

To explore if the effects of these risk factors were consistent across breeds, the same primary risk factor model was fit to a subset of breeds. The top three most frequent breeds within each weight quartile were selected as an analysis subset (see **Statistical Methods: Breed-Specific Analyses** section for more information). **Supplemental Table 1, Supplemental Figure 2** and **Supplemental Figure 3** the 12 representative breeds as well as breed-specific OA incidence tested against the overall cohort OA incidence rate.

**Figure 2** summarizes these results using a dot-and-whisker plot of the standardized HR and 95% CI for each of the risk factors within each breed. Higher weight and older age are consistently associated with higher risks of OA across breeds. However, the effect of desexing on OA incidence shows variability across breeds, where desexing does not appear to be a risk factor for Yorkshire Terriers, Maltese, Shih Tzus, Pugs, Dachshunds and Beagles. The effects of BCS also differ across breeds, where effects are more pronounced in smaller breeds. HRs and 95% CIs reported in **Supplemental Table 2.**

**Figure 2.**
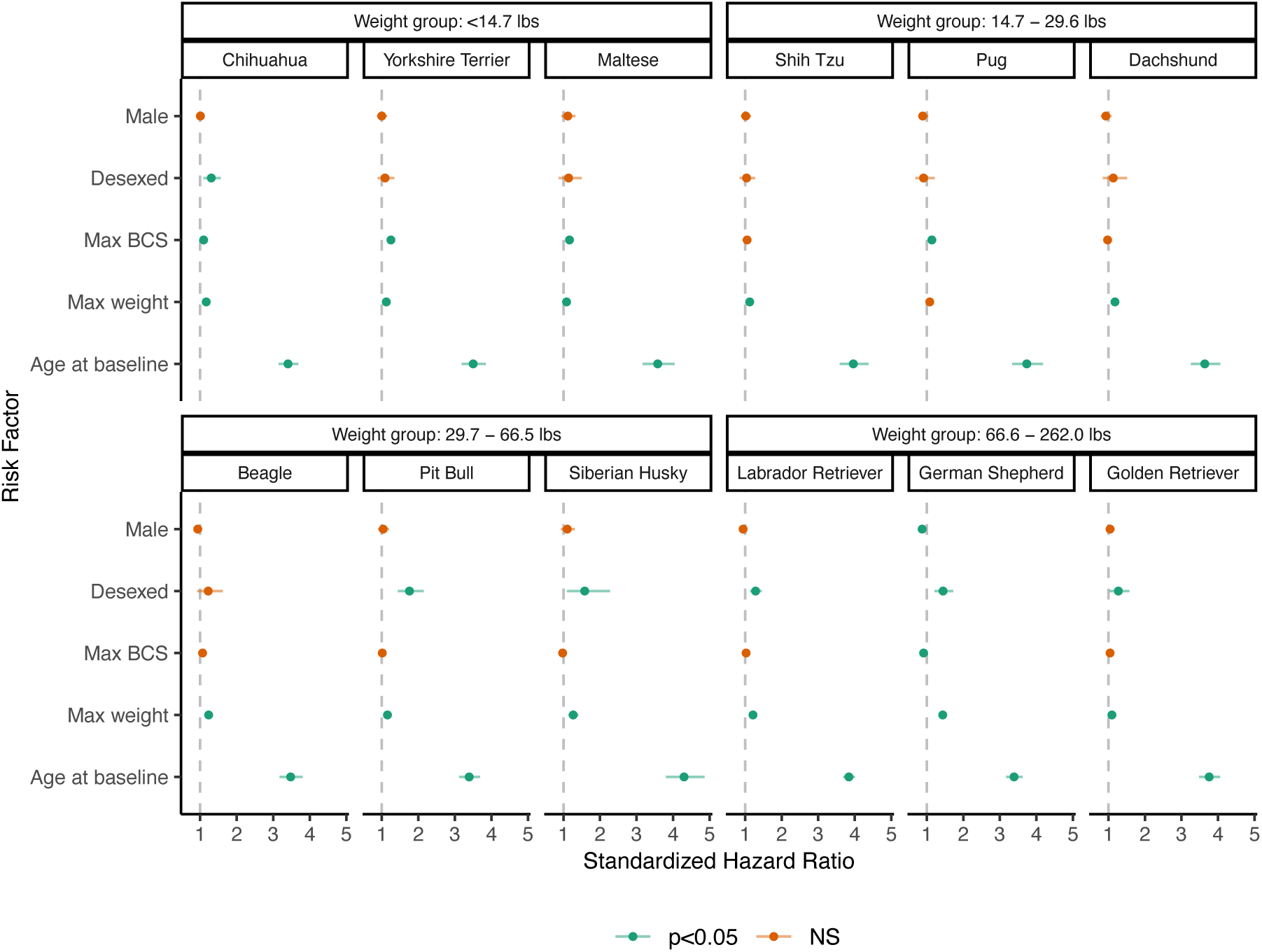
Dot-and-whisker plot results of breed-specific analysis of primary risk factors of OA. Dots and whiskers reflect the standardized HR and 95% CIs associated with each risk factor, respectively. Each panel reflects the model fit to an individual breed, where panels are grouped by weight groups. Color assignments designate statistical significance of the HR.

### Does desexing moderate the effect of max body weight on OA?

Both weight and desexing are independent risk factors of OA. However, it has been reported that dogs who are desexed may be more likely to develop obesity,(21,22,37) which may increase the risk of OA. To test if the influence of higher body weight on OA-free survival differed by desexing status, the interaction between adult weight and desexing status was tested.

There was a statistically significant interaction between desexing and adult weight (Std. HR=1.07 (1.03, 1.11), **Table 4**), suggesting that the increased risk of OA in larger dogs is greater in those that are desexed compared to intact dogs. **Figure 3** shows the predicted survival curves as well as the predicted median survival times for each weight class and desexing group. Both desexed and intact dogs show that higher body weight is associated with higher OA risk; however, this risk appears slightly larger in desexed dogs compared to intact dogs.

**Figure 3.**
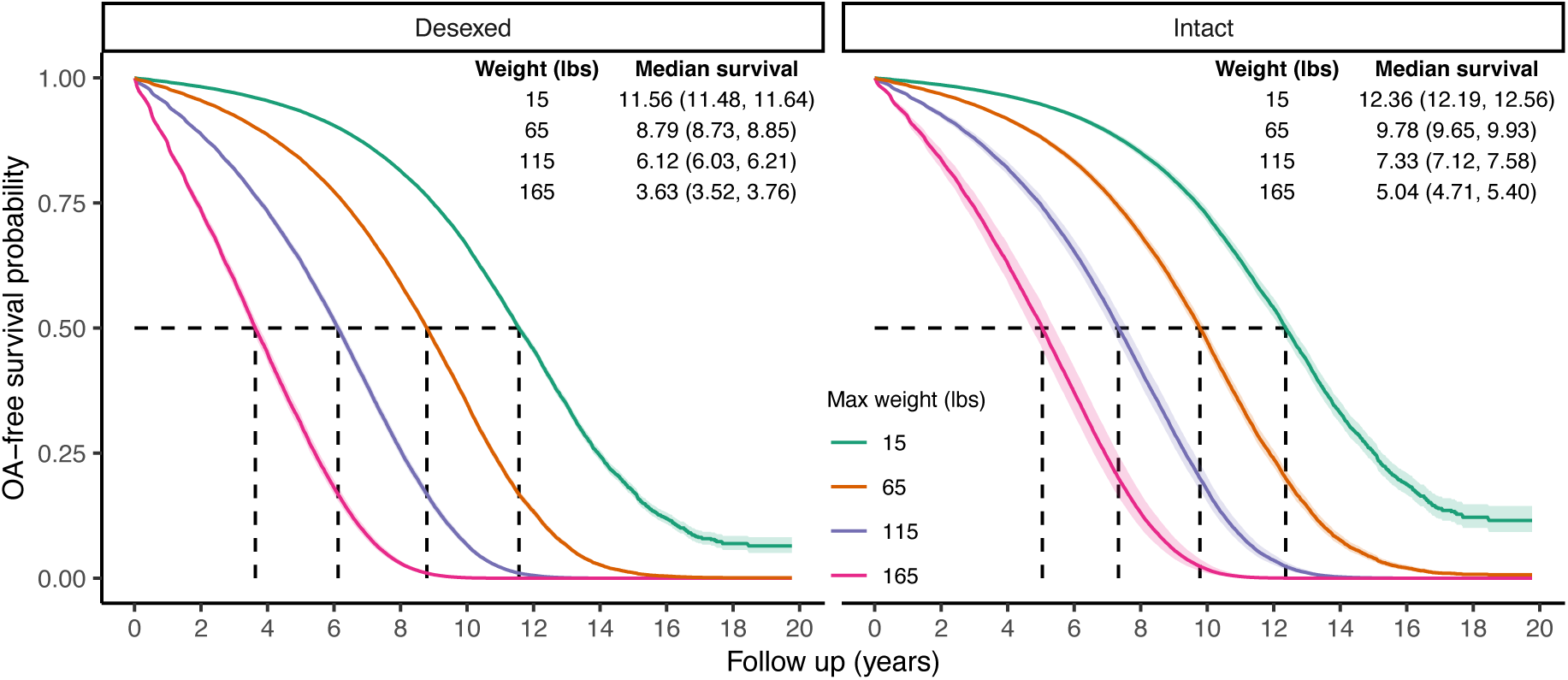
Predicted marginal OA-free survival curves to visualize interaction effects between weight and desexing status. Color indicates maximum weight values. Median survival times and 95% CIs are reported. To generate predictions, median values were used for continuous covariates, proportions for binary variables, and the presence of wellness plan were used.

**Table 4.** Stratified Cox proportional hazard model testing the interaction between desexing and maximum adult weight and BCS.

| Model | Variable | HR (95% CI) | Std. HR (95% CI) | p-value | Concordance | N obs |
| --- | --- | --- | --- | --- | --- | --- |
| Desexing x Weight | Age at baseline | 1.35 (1.35-1.36) | 3.70 (3.64-3.76) | <0.001*** | 0.79 | 121,035 |
|  | Max weight | 1.02 (1.02-1.02) | 1.73 (1.67-1.79) | <0.001*** |  |  |
|  | Max BCS | 1.02 (1.01-1.03) | 1.02 (1.01-1.03) | <0.001*** |  |  |
|  | Desexed | 1.23 (1.15-1.32) | 1.34 (1.29-1.40) | <0.001*** |  |  |
|  | Male | 0.91 (0.89-0.93) | 0.91 (0.89-0.93) | <0.001*** |  |  |
|  | Desexed x max weight | 1.00 (1.00-1.00) | 1.07 (1.03-1.11) | <0.001*** |  |  |
| Desexing x BCS | Age at baseline | 1.35 (1.35-1.36) | 3.70 (3.64-3.76) | <0.001*** | 0.79 | 121,035 |
|  | Max weight | 1.02 (1.02-1.02) | 1.83 (1.81-1.85) | <0.001*** |  |  |
|  | Max BCS | 1.02 (0.99-1.06) | 1.03 (0.99-1.07) | 0.186 |  |  |
|  | Desexed | 1.42 (1.14-1.76) | 1.37 (1.31-1.43) | 0.002** |  |  |
|  | Male | 0.91 (0.89-0.93) | 0.91 (0.89-0.93) | <0.001*** |  |  |
|  | Desexed x max BCS | 0.99 (0.96-1.03) | 0.99 (0.95-1.04) | 0.754 |  |  |

\*Std. HR= Standardized continuous variables (mean centered, scaled by standard deviation). Wellness plan treated as stratifying variable.

### Does weight gain increase risk of developing OA?

Limitations of the modeling approach used in **Table 2** and **Figure 1** include the lack of temporal precedence – specifically, the inability to test subsequent weight gain after desexing and its influence on OA incidence. To test this hypothesis, percent changes in maximum weight within desexed dogs (calculated as 100*[maximum weight - mature weight]/mature weight) was tested as a risk factor for OA.

Both main effects of percent weight change as well as the interaction with mature body weight were tested in the full cohort and in a sensitivity analysis restricting to dogs whose age at desexing was ≤ 2 years of age. Mature body weight was treated as a covariate as larger-breed dogs are less likely to have larger percent changes in body weight compared to smaller-breed dogs. To determine if weight changes were independent of desexing status, these analyses were performed in intact dogs as well, where percent changes in weight reflected overall changes from mature body weight to maximum body weight.

Results in **Table 5** show that positive percent change in body weight was associated with higher risk of OA (full cohort Std. HR=1.12 (1.09-1.15); age desexing ≤ 2 Std. HR=1.12 (1.09- 1.15)). There was also a significant interaction effect between percent change in body weight after desexing and mature body size (Std. HR=1.09 (1.05-1.12)), which held in the dogs desexed ≤ 2 years of age. These results suggest that percent increases in body weight after desexing may have stronger deleterious effects in larger dogs, compared to smaller dogs.

**Table 5.** Stratified Cox proportional hazard model testing both the main effects and interaction effects of percent change in body weight in desexed dogs, dogs desexed ≤ 2 years of age, and intact dogs, respectively. For dogs who were desexed, percent change in body weight is based on weight change exclusively after desexing.

| Subset | Model | Variable | HR (95% CI) | Std. HR (95% CI) | p-value | Concordance | N obs |
| --- | --- | --- | --- | --- | --- | --- | --- |
| All desexed dogs | Main effect | Age at baseline | 1.20 (1.15-1.26) | 2.23 (1.82-2.74) | <0.001*** | 0.69 | 23,099 |
|  |  | Male | 0.86 (0.82-0.90) | 0.86 (0.82-0.90) | <0.001*** |  |  |
|  |  | % change in weight after desexing | 1.01 (1.01-1.01) | 1.12 (1.09-1.15) | <0.001*** |  |  |
|  |  | Mature weight | 1.02 (1.02-1.02) | 1.95 (1.91-2.00) | <0.001*** |  |  |
|  | Interaction | Age at baseline | 1.21 (1.15-1.26) | 2.25 (1.83-2.76) | <0.001*** | 0.69 | 23,099 |
|  |  | Male | 0.86 (0.82-0.91) | 0.86 (0.82-0.91) | <0.001*** |  |  |
|  |  | % change in weight after desexing | 1.00 (1.00-1.01) | 1.05 (1.01-1.09) | 0.027* |  |  |
|  |  | Mature weight | 1.02 (1.02-1.02) | 1.86 (1.80-1.92) | <0.001*** |  |  |
|  |  | % change in weight after desexing x mature weight | 1.00 (1.00-1.00) | 1.09 (1.05-1.12) | <0.001*** |  |  |
| Dogs desexed $\leq 2$ years old | Main effect | Age at baseline | 1.12 (0.98-1.28) | 1.02 (1.00-1.05) | 0.108 | 0.69 | 21,296 |
|  |  | Male | 0.86 (0.82-0.90) | 0.86 (0.82-0.90) | <0.001*** |  |  |
|  |  | % change in weight after desexing | 1.01 (1.01-1.01) | 1.12 (1.09-1.15) | <0.001*** |  |  |
|  |  | Mature weight | 1.02 (1.02-1.02) | 2.00 (1.95-2.05) | <0.001*** |  |  |
|  | Interaction | Age at baseline | 1.12 (0.98-1.28) | 1.02 (1.00-1.05) | 0.097 | 0.69 | 21,296 |
|  |  | Male | 0.86 (0.82-0.91) | 0.86 (0.82-0.91) | <0.001*** |  |  |
|  |  | % change in weight after desexing | 1.00 (1.00-1.01) | 1.04 (1.00-1.08) | 0.057 |  |  |
|  |  | Mature weight | 1.02 (1.02-1.02) | 1.89 (1.83-1.96) | <0.001*** |  |  |
|  |  | % change in weight after desexing x mature weight | 1.00 (1.00-1.00) | 1.09 (1.05-1.13) | <0.001*** |  |  |
| Intact Dogs | Main effect | Age at baseline | 1.44 (1.30-1.59) | 4.84 (3.12-7.50) | <0.001*** | 0.69 | 5,519 |
|  |  | Male | 1.23 (1.04-1.44) | 1.23 (1.04-1.44) | 0.014* |  |  |
|  |  | % change in weight | 1.01 (1.01-1.01) | 1.19 (1.14-1.25) | <0.001*** |  |  |
|  |  | Mature weight | 1.02 (1.02-1.02) | 1.70 (1.59-1.81) | <0.001*** |  |  |
|  | Interaction | Age at baseline | 1.45 (1.31-1.60) | 4.97 (3.20-7.71) | <0.001*** | 0.69 | 5,519 |
|  |  | Male | 1.26 (1.07-1.48) | 1.26 (1.07-1.48) | 0.006** |  |  |
|  |  | % change in weight | 1.01 (1.00-1.01) | 1.12 (1.05-1.20) | 0.001** |  |  |
|  |  | Mature weight | 1.01 (1.01-1.02) | 1.57 (1.45-1.71) | <0.001*** |  |  |
|  |  | % change in weight x mature weight | 1.00 (1.00-1.00) | 1.13 (1.05-1.22) | 0.001** |  |  |

\*Std. HR= Standardized continuous variables (mean centered, scaled by standard deviation). Wellness plan treated as stratifying variable.

In order to illustrate these effects in more interpretable units, **Figure 4** shows the main effects of percent changes in body weight after desexing, where every 50% increase in body weight after desexing was associated with an approximately 1 year decrease in median time to OA. **Figure 5** visualizes the finding that percent increases in body weight after desexing was associated with higher OA risks in larger dogs compared to smaller dogs. It is important to note that 50% increases in body weight become less likely as mature body size increases and the use of 50% increments are used only to facilitate interpretation of the direction and magnitude of effects.

**Figure 4.**
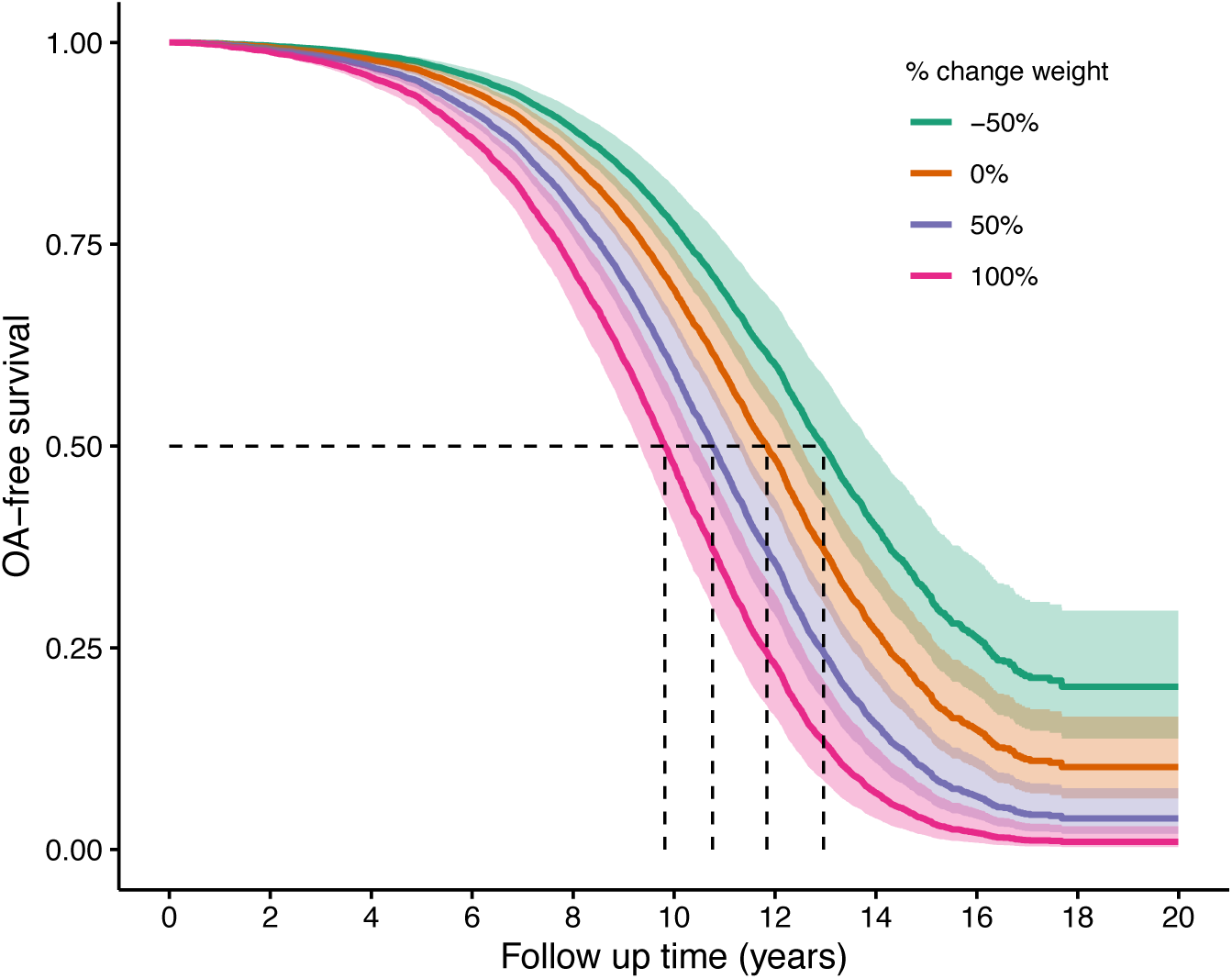
Predicted marginal OA-free survival curves illustrating the main effect of percent weight changes after desexing. Color indicates percent change values. To generate predictions, median values were used for continuous covariates, proportions for binary variables, and the presence of wellness plan were used.

**Figure 5.**
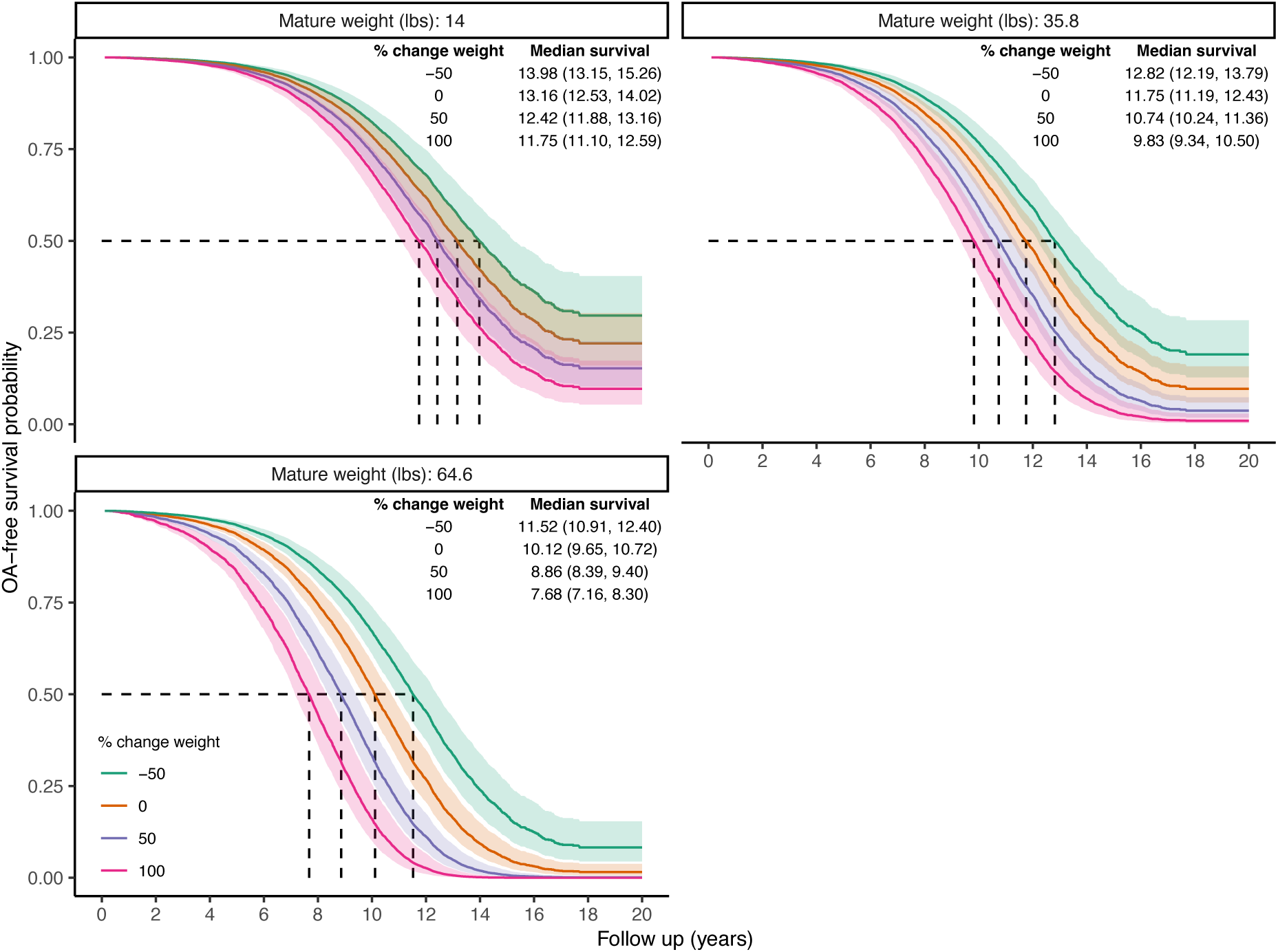
Predicted marginal OA-free survival curves illustrating the interaction effect between percent weight change after desexing and mature body weight. Color indicates percent change values. Median survival times and 95% CIs are reported. To generate predictions, median values were used for continuous covariates, proportions for binary variables, and the presence of wellness plan were used.

### Does increased BCS influence OA risks?

As BCS is a distinct construct from weight or body size, percent changes in body condition score after desexing was also tested as a risk factor for OA. Paralleling the approach used in the percent change in weight models, both main effects of percent BCS change as well as the interaction with mature BCS were tested in the full cohort and in a sensitivity analysis restricting to dogs whose age at desexing was ≤ 2 years of age. Mature body BCS was treated as a covariate as dogs with higher BCS at maturity are less likely to have larger BCS changes in BCS compared to dogs with lower BCS at maturity. To determine if weight changes were independent of desexing status, these analyses were performed in intact dogs as well, where percent changes in BCS reflected overall changes from mature BCS to maximum BCS.

Results indicate increases in percent change in BCS are associated with higher risk of OA (Std. HR=1.12 (1.05-1.20), **Table 6**), and that these effects may be stronger in dogs with higher mature BCS (Std. HR=1.16 (1.04-1.29), **Table 6**). However, limited variability in BCS in this cohort makes determining the clinical significance of these effects difficult. As seen in **Figure 6** and **Figure 7**, standard errors from model predictions show significant overlap, except in dogs who have higher mature BCS, where effects begin to appear strongest. However, it is important to note that due to there being a ceiling in the BCS tool, dogs with mature BCS of 7 can only experience at most a percent change increase of 28% (28%=100*(9-7)/7).

**Figure 6.**
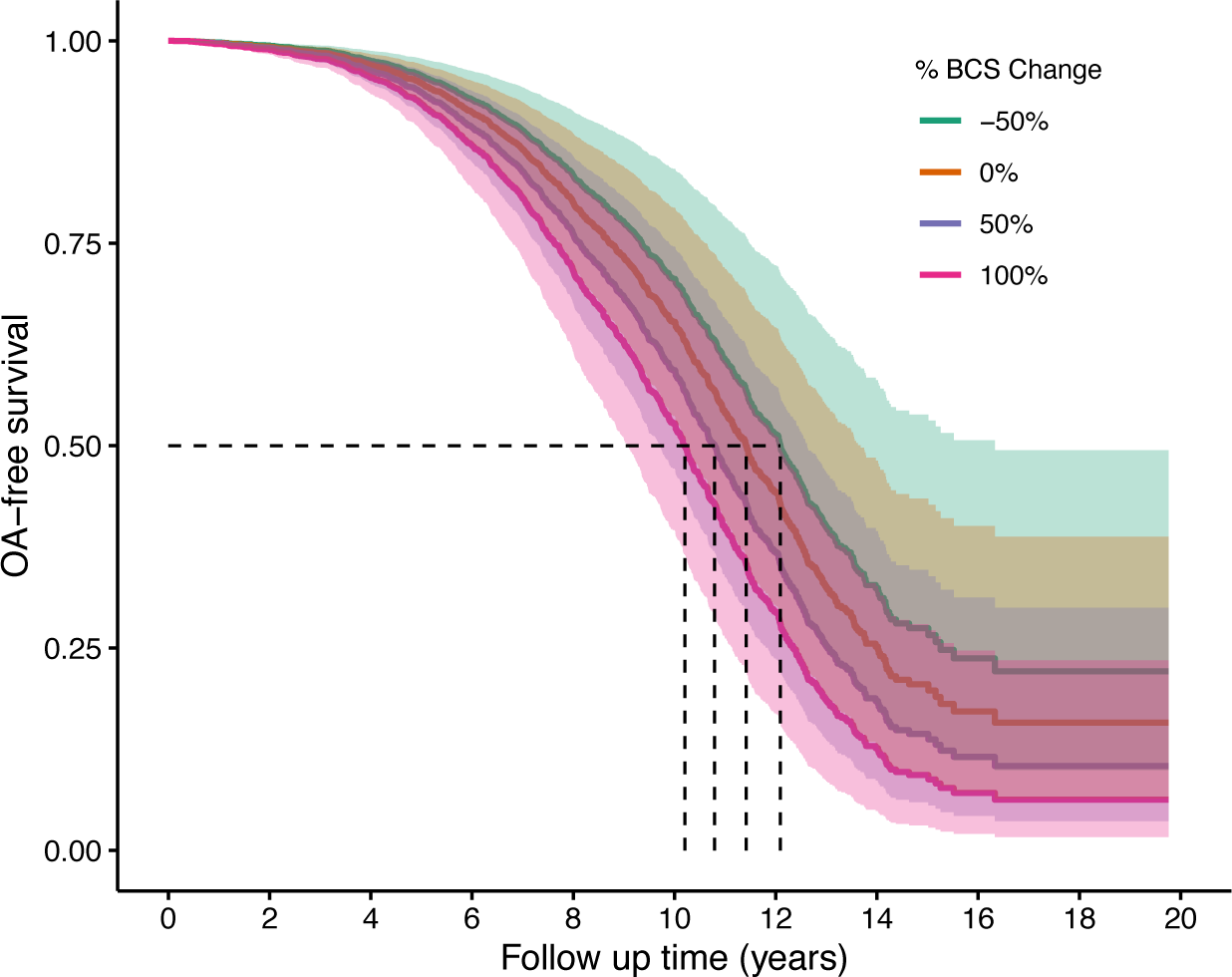
Predicted marginal OA-free survival curves illustrating the main effect of percent BCS after desexing. Color indicates percent change values. To generate predictions, median values were used for continuous covariates, proportions for binary variables, and the presence of wellness plan were used.

**Figure 7.**
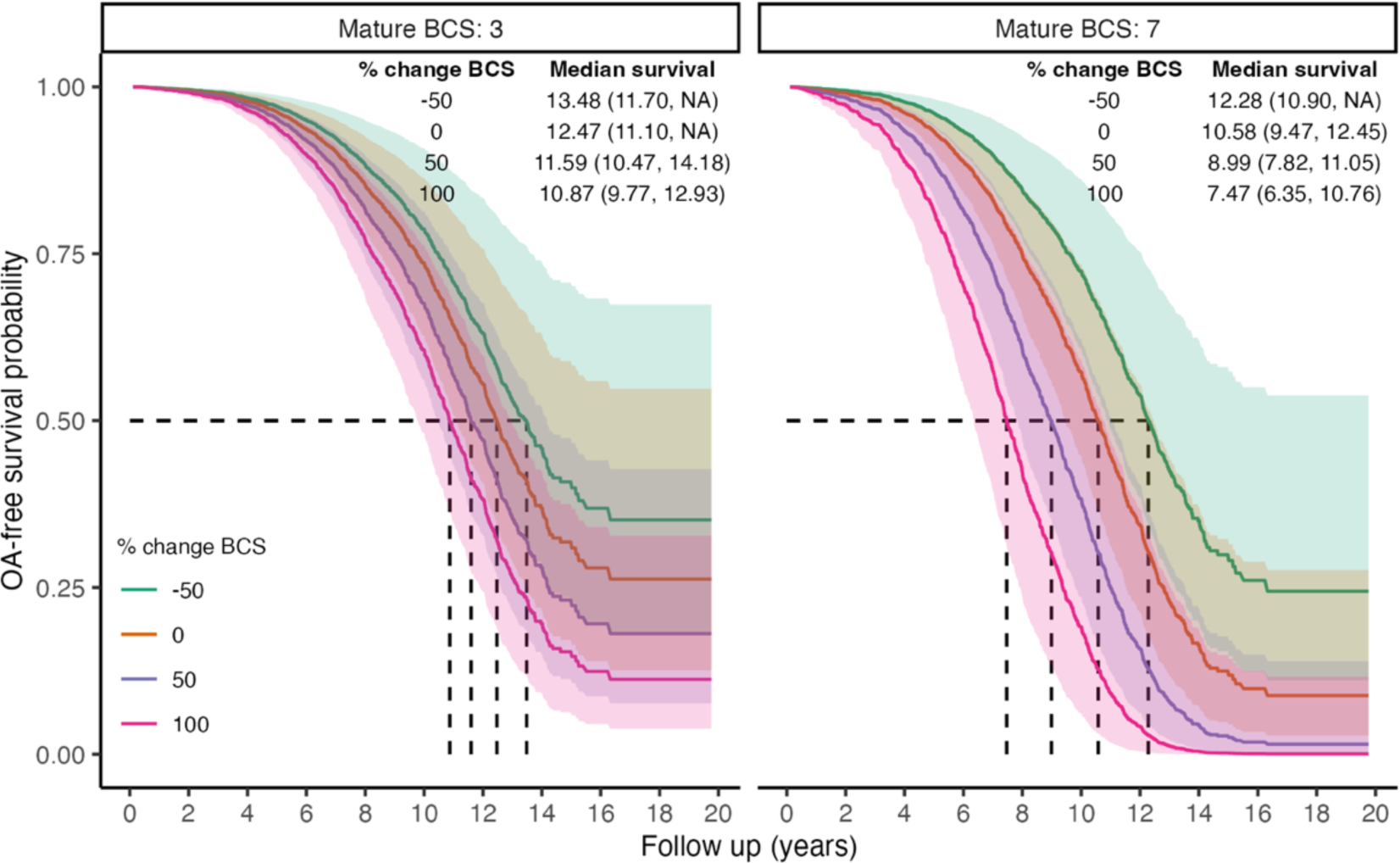
Predicted marginal OA-free survival curves illustrating the interaction effect between percent BCS after desexing and mature BCS. Color indicates percent change values. Median survival times and 95% CIs are reported. To generate predictions, median values were used for continuous covariates, proportions for binary variables, and the presence of wellness plan were used.

**Table 6.** Stratified Cox proportional hazard model testing both the main effects and interaction effects of percent change in BCS in desexed dogs, dogs desexed ≤ 2 years of age, and intact dogs, respectively. For dogs who were desexed, percent change in BCS is based on BCS change exclusively after desexing.

| Subset | Model | Variable | HR (95% CI) | Std. HR (95% CI) | p-value | Concordance | N obs |
| --- | --- | --- | --- | --- | --- | --- | --- |
| All desexed dogs | Main effect | Age at baseline | 1.25 (1.12-1.39) | 2.60 (1.64-4.12) | <0.001*** | 0.67 | 7,484 |
|  |  | Male | 0.92 (0.82-1.04) | 0.92 (0.82-1.04) | 0.186 |  |  |
|  |  | % change in BCS after desexing | 1.00 (1.00-1.01) | 1.12 (1.05-1.20) | <0.001*** |  |  |
|  |  | Mature weight | 1.02 (1.02-1.02) | 1.69 (1.61-1.78) | <0.001*** |  |  |
|  |  | Mature BCS | 1.16 (1.08-1.24) | 1.18 (1.09-1.29) | <0.001*** |  |  |
|  | Interaction | Age at baseline | 1.24 (1.12-1.38) | 2.56 (1.61-4.05) | <0.001*** | 0.67 | 7,484 |
|  |  | Male | 0.92 (0.82-1.04) | 0.92 (0.82-1.04) | 0.166 |  |  |
|  |  | % change in BCS after desexing | 1.00 (1.00-1.00) | 1.01 (0.90-1.13) | 0.831 |  |  |
|  |  | Mature weight | 1.02 (1.02-1.02) | 1.69 (1.60-1.78) | <0.001*** |  |  |
|  |  | Mature BCS | 1.16 (1.08-1.25) | 1.19 (1.09-1.29) | <0.001*** |  |  |
|  |  | % change in BCS after desexing x mature BCS | 1.00 (1.00-1.00) | 1.16 (1.04-1.29) | 0.008** |  |  |
| Dogs desexed $\leq 2$ years old | Main effect | Age at baseline | 1.16 (0.80-1.66) | 1.03 (0.96-1.11) | 0.432 | 0.67 | 6,921 |
|  |  | Male | 0.91 (0.81-1.03) | 0.91 (0.81-1.03) | 0.135 |  |  |
|  |  | % change in BCS after desexing | 1.00 (1.00-1.01) | 1.14 (1.06-1.21) | <0.001*** |  |  |
|  |  | Mature weight | 1.02 (1.02-1.02) | 1.72 (1.63-1.81) | <0.001*** |  |  |
|  |  | Mature BCS | 1.15 (1.07-1.24) | 1.17 (1.08-1.27) | <0.001*** |  |  |
|  | Interaction | Age at baseline | 1.14 (0.80-1.64) | 1.03 (0.95-1.11) | 0.471 | 0.67 | 6,921 |
|  |  | Male | 0.91 (0.81-1.03) | 0.91 (0.81-1.03) | 0.124 |  |  |
|  |  | % change in BCS after desexing | 1.00 (1.00-1.01) | 1.03 (0.92-1.14) | 0.647 |  |  |
|  |  | Mature weight | 1.02 (1.02-1.02) | 1.71 (1.62-1.81) | <0.001*** |  |  |
|  |  | Mature BCS | 1.16 (1.07-1.25) | 1.18 (1.08-1.28) | <0.001*** |  |  |
|  |  | % change in BCS after desexing x mature BCS | 1.00 (1.00-1.00) | 1.16 (1.05-1.29) | 0.005** |  |  |
| Intact dogs | Main effect | Age at baseline | 2.07 (1.52-2.82) | 23.46 (6.15-89.40) | <0.001*** | 0.67 | 2,153 |
|  |  | Male | 1.21 (0.73-2.01) | 1.21 (0.73-2.01) | 0.467 |  |  |
|  |  | % change in BCS | 1.00 (1.00-1.01) | 1.13 (0.98-1.30) | 0.085 |  |  |
|  |  | Mature weight | 1.02 (1.01-1.02) | 1.67 (1.39-2.00) | <0.001*** |  |  |
|  |  | Mature BCS | 1.00 (0.82-1.21) | 1.00 (0.80-1.25) | 0.981 |  |  |
|  | Interaction | Age at baseline | 2.07 (1.51-2.82) | 23.16 (6.02-89.13) | <0.001*** | 0.67 | 2,153 |
|  |  | Male | 1.21 (0.73-2.02) | 1.21 (0.73-2.02) | 0.463 |  |  |
|  |  | % change in BCS | 1.00 (1.00-1.01) | 1.12 (0.92-1.36) | 0.267 |  |  |
|  |  | Mature weight | 1.02 (1.01-1.02) | 1.66 (1.39-2.00) | <0.001*** |  |  |
|  |  | Mature BCS | 1.00 (0.81-1.24) | 1.00 (0.79-1.28) | 0.971 |  |  |
|  |  | % change in BCS x mature BCS | 1.00 (1.00-1.00) | 1.02 (0.76-1.39) | 0.876 |  |  |

\*Std. HR= Standardized continuous variables (mean centered, scaled by standard deviation).
Wellness plan treated as stratifying variable.

These analyses were also performed in intact dogs, where percent changes in BCS reflect overall changes from mature BCS to maximum BCS to identify any changes across desexing status, where percent changes in BCS were not statistically significantly associated with OA incidence.

### Does age at desexing influence OA risks?

To test the hypothesis that age at desexing is a risk factor for OA, age at desexing was added into the primary risk factor model. Age at desexing was tested as a continuous variable as well as a categorical variable (<6 months, 6-12 months, 12-24 months, and 24-60 months). Age at desexing was significantly inversely associated with OA risk (Std. HR=0.90 (0.86-0.94) **Table 7**). Categorical analysis shows that relative to dogs desexed before 6 months of age, dogs desexed 6-12 months are 13% less likely to develop OA, while dogs who are desexed 12-24 or 24-60 months are 28% and 32% less likely to develop OA, respectively. This categorical model suggests a potential non-linear relationship between age at desexing and OA risk, such that the effects of age at desexing diminish as dogs get older. To test this, a sensitivity analysis was performed adding a quadratic term in the continuous age at desexing model. The quadratic term for age at desexing was statistically significant (Std. HR=1.35 (1.22-1.49) **Table 7**)). Predictions from all models are presented in **Figure 8**. Results from this quadratic model more closely relate to the categorical model, providing additional evidence that the magnitude of the OA-related risk associated with age at desexing diminishes with age.

**Figure 8.**
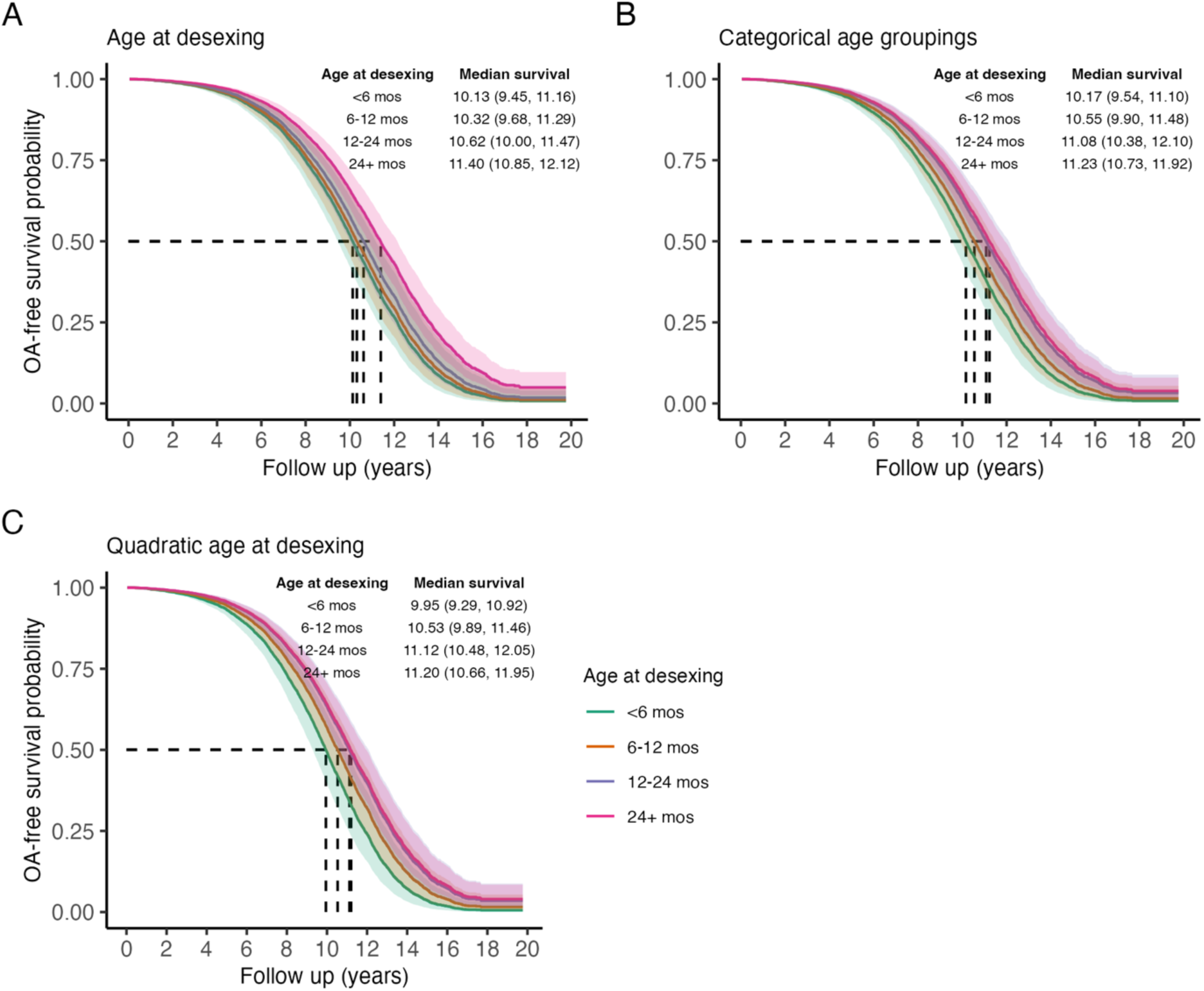
Predicted marginal OA-free survival curves illustrating the effect of age at desexing on OA incidence, treating age as a linear continuous variable, a categorical variable, or quadratic variable. Color indicates ages at desexing. Median survival times and 95% CIs are reported. To generate predictions, median values were used for continuous covariates, proportions for binary variables, and the presence of wellness plan were used.

**Table 7.** Stratified Cox proportional hazard model testing the role of age at desexing as a risk factor for OA. Age at desexing is treated as a continuous measure and as a categorical variable.

| Desexing variable | Variable | HR (95% CI) | Std. HR (95% CI) | p-value | Concordance | N obs |
| --- | --- | --- | --- | --- | --- | --- |
| Continuous | Age at baseline | 1.42 (1.32-1.53) | 1.42 (1.32-1.53) | <0.001*** | 0.70 | 24,159 |
|  | Max weight | 1.02 (1.02-1.02) | 1.83 (1.79-1.87) | <0.001*** |  |  |
|  | Max BCS | 1.05 (1.02-1.07) | 1.05 (1.03-1.08) | <0.001*** |  |  |
|  | Male | 0.88 (0.84-0.93) | 0.88 (0.84-0.93) | <0.001*** |  |  |
|  | Age at desexing | 0.87 (0.82-0.92) | 0.90 (0.86-0.94) | <0.001*** |  |  |
| Quadratic | Age at baseline | 1.42 (1.32-1.52) | 1.42 (1.32-1.52) | <0.001*** | 0.70 | 24,159 |
|  | Max weight | 1.02 (1.02-1.02) | 1.82 (1.79-1.86) | <0.001*** |  |  |
|  | Max BCS | 1.05 (1.02-1.07) | 1.06 (1.03-1.08) | <0.001*** |  |  |
|  | Male | 0.89 (0.84-0.93) | 0.89 (0.84-0.93) | <0.001*** |  |  |
|  | Age at desexing | 0.61 (0.53-0.69) | 0.67 (0.61-0.75) | <0.001*** |  |  |
|  | Age at desexing<sup>2</sup> | 1.09 (1.06-1.13) | 1.35 (1.22-1.49) | <0.001*** |  |  |
| Grouped | Age at baseline | 1.41 (1.31-1.51) | 4.39 (3.26-5.90) | <0.001*** | 0.70 | 24,159 |
|  | Max weight | 1.02 (1.02-1.02) | 1.82 (1.78-1.86) | <0.001*** |  |  |
|  | Max BCS | 1.05 (1.02-1.07) | 1.06 (1.03-1.08) | <0.001*** |  |  |
|  | Male | 0.89 (0.85-0.93) | 0.89 (0.85-0.93) | <0.001*** |  |  |
|  | Desexed <6 mos | Reference |  |  |  |  |
|  | Desexed 6-12 mos | 0.87 (0.82-0.92) | 0.87 (0.82-0.92) | <0.001*** |  |  |
|  | Desexed 12-24 mos | 0.72 (0.64-0.80) | 0.72 (0.64-0.80) | <0.001*** |  |  |
|  | Desexed 24-60 mos | 0.68 (0.58-0.79) | 0.68 (0.58-0.79) | <0.001*** |  |  |

\*Std. HR= Standardized continuous variables (mean centered, scaled by standard deviation). Wellness plan treated as stratifying variable.

## Discussion

The proportion of dogs in this population that developed OA was 24.9%. It is difficult to compare this to occurrence data from other studies due to differences in study populations, case definitions, and other methodological factors. Previous analyses of primary practice databases in the UK, for example, reported a one-year period prevalence of OA in dogs of any age of 2.34%(1) and 2.5%.(4) An owner survey, also in the UK, reported an overall prevalence of 2.23%.(38)

In contrast, a proprietary industry survey of 200 veterinarians in the U.S. reported an overall prevalence of 20%(5) in dogs over 1 year of age, a figure frequently cited in reviews and other articles discussing the condition. A more recent study involving dogs 5-12 years of age and over 11 kg in weight presenting for dental prophylaxis found 68% of dogs had radiographic evidence of OA in one or more joints.(39) However, owner surveys from a subset of the population reported no clinical signs in 71% of the dogs, and these would likely not have received a diagnosis of OA in routine clinical practice. Such variability creates significant uncertainty about the true rate of occurrence of OA in dogs.

In this study, the incidence of 24.9% represents the clinical diagnosis of OA at any age and by any criteria in dogs from a primary care population who died in 2019, with no fixed interval or duration of follow up. Diagnoses could be based on clinical impression, history and physical examination, radiographic interpretations, or other diagnostic procedures. Given the large size and broad geographic distribution of the study population, this should be reasonably representative of the occurrence of OA as routinely identified by general practice clinicians in companion dogs in the U.S.

Detection bias in retrospective cohort studies can influence both the apparent prevalence of a condition and the apparent impact of risk factors for the disease.(40) Owners of dogs in this population were offered various wellness plans, which consist of a prepaid packages of services that can reduce the cost of clinic visits and some diagnostic tests, including radiographs. Enrollment in such a wellness plan has previously been suggested to be a risk factor for detection bias,(37) and in this study it was strongly associated with OA diagnosis. Dogs with a wellness plan had far more clinic visits and were more likely to have radiographs than dogs not enrolled in such a plan (data not shown), likely leading to a higher rate of detection of OA. This factor was controlled for by stratification in the analysis of other putative risk factors.

Age was by far the strongest risk factor for OA diagnosis, consistent with the established understanding of OA as a disease of aging. This effect was consistent across breeds of different sizes and with different rates of OA occurrence (**Figure 2** and **Supplemental Table 2**). Diagnosis of OA in individual patients is more likely as time passes, both because it is an incurable condition and so will be present and available for diagnosis longer, and because it is progressive and so more likely to become clinically apparent as a dog ages.

In previous studies, sex has been inconsistently identified as a risk factor for OA.(6) In this study, males were at lower risk, though the size of this effect in the overall population was small. Within breeds, males were at lower risk in some breeds but not others, independent of body weight and BCS (**Figure 2** and **Supplemental Table 2**).

All types of OA were included in the case definition for this study, regardless of which joint was affected or whether a predisposing condition, such as joint dysplasia or cranial cruciate ligament rupture, was identified. Previous evaluations of sex as a risk factor for OA have focused more narrowly on particular joints, predisposing conditions, or breeds, and it is likely that the role of sex as a risk factor may vary among these.(4) The small overall risk difference between males and females suggests that diagnostic and preventative measures should be directed equally at dogs of both sexes.

Previous studies have shown that OA risk differs markedly between breeds.(6) Breed is a complex risk factor involving differences in genetic makeup, body size and conformation, and likely also lifestyle variables influenced by owners, such as feeding practices and the type and intensity of activity. Because of the enormous phenotypic variability among dogs, evaluation of the role of body weight in OA risk can easily confound the effects of body size and breed with those of overweight.

In general, larger body size is often identified as a risk factor for OA in dogs. This pattern was confirmed in this study, with increased mature body weight positively associated with OA risk. When OA risk was compared among breeds, larger breeds had higher risk, consistent with previous findings **(Supplemental Figures 2** and **3** and **Supplemental Table 2**).

In an effort to disentangle the effects of body size and overweight to some extent, we examined the interaction between the percentage increase above mature body weight and the risk for OA. This analysis showed that weight gain after maturity, likely representing excess adipose mass, increases risk of OA. This effect is greater in larger dogs, possibly due to an exacerbation of the already elevated risk posed by larger body size, or because the absolute mechanical burden of weight gained after maturity is greater in larger dogs.

Ideally, body condition score should help to isolate overweight and obesity as a risk factor independent of mature body size, so it was expected that BCS would be a strong predictor of OA risk. In this study population, 40.1% of dogs were classed as overweight (BCS 6-7/9) and 4.1% as obese (BCS 8-9/9), and the proportion of overweight and obesity was higher in the dogs that developed OA (overweight= 50.8%, obese= 5.3%) than in those that did not (overweight= 36.7%, obese= 3.8%). These figures are broadly consistent with previous reports, though the estimated prevalence of overweight and obesity vary widely depending on the population, time period, assessment methods, and category definitions.(21,41–43)

While BCS was significantly correlated with OA risk, the size of this effect was very small. This relationship was stronger in smaller dogs, unlike the effect of weight gain after maturity. Overall, BCS was not a powerful predictor of OA diagnosis. A possible explanation for this unexpected result may be practical challenges in the implementation of BCS scoring. There are several scoring systems in use by veterinarians, and the validation data for these are limited, involving relatively small numbers of dogs, breeds, and clinicians.(44–47) Some studies have reported that BCS scoring correlates well with objective measures of body fat.(46) However, the strength of this correlation varies significantly between breeds. The consistency and reliability of BCS scoring among veterinarians scoring dogs of many different breeds and conformations in general practice is uncertain.

Various assessment tools for body condition were used at different times in evaluation of this population, and scores from a 3-point and a 5-point scale were mapped onto a 9-point scale for analysis. In principle, a system with more gradations should allow for a more accurate distinction between degrees of overweight among dogs and help to identify more clearly the relationship between BCS and clinical conditions such as OA. In practice, however, the distribution of BCS scores in this data set was clearly discontinuous, with the vast majority of dogs scored as 5/9 (45.4%) or 7/9 (35%). This suggests clinicians tended to lump dogs into broad “normal” and “overweight” categories, rather than utilizing BCS scoring as a more finely graded, continuous measure.

Though this was true across all sizes of dogs, there was greater use of the full range of scores, particularly those below 5/9, in smaller dogs (**Supplemental Figure 4**). This suggests clinicians may find it easier to distinguish gradations of body condition in smaller breeds. The fact that BCS was a stronger predictor of OA in smaller dogs, among which BCS scores were more granular, supports the possibility that the lumping of many dogs into either “normal” or “overweight” obscured the overall relationship between BCS and OA.

Further evaluation of the reliability of extant BCS scoring systems in heterogenous general practice populations, and the development and validation of alternative assessment tools that obviate the limitations of BCS, would be useful in clarifying the role of overweight and obesity as risk factors for disease.

Neuter status has been consistently identified as a risk for OA, with intact individuals at lower risk than neutered dogs. There is also evidence that neutering before 12 months of age is associated with an increased risk of several of the most important conditions predisposing to OA (hip and elbow dysplasia and cranial cruciate ligament rupture), though this effect may only appear in larger dogs (over approximately 43 lbs in one recent analysis).(18)

This study is consistent with previous findings that the risk of OA is increased following desexing in most medium and large breed dogs (over 30 lbs) and in some smaller breeds. While predisposing conditions were not evaluated, the impact of neutering on OA risk overall increased with body size, and neutering was less frequently associated with OA risk in smaller breeds, supporting the apparent greater significance of neutering as a risk factor in larger dogs.

Neutering is also an established risk factor for obesity,(21,22,37) and it is possible that one way in which neutering increases OA risk is by increasing the propensity for overweight and obesity. Because of challenges in the interpretation of BCS scores, it was difficult to compare the occurrence of overweight and obesity between intact and neutered dogs. The effect of body weight gain after maturity on OA risk was not different between intact and neutered dogs, so overweight and obesity are clearly important risk factors in all dogs regardless of neuter status. While increasing the incidence of overweight and obesity may be one mediator of the impact of neutering on OA risk, this hypothesis could not be directly evaluated in this study.

In this study, age at neutering was inversely associated with OA risk in the total population, indicating that risk decreased progressively with delayed neutering up to 2 years of age. This effect was the same in dogs of different body sizes, which is not consistent with previous reports suggesting earlier neutering (before 12 months of age) increased the risk of predisposing conditions, such as hip and elbow dysplasia and cruciate ligament rupture, in larger dogs more than smaller dogs.(18) In this large a cohort, the power to detect such a difference in risk associated with size should be high, so these data suggest the OA risk associated with age at neutering would apply to all dogs in similar, heterogenous primary care populations. Differences between this study and others in the appearance of a size effect for this risk factor may be explained by differences in detection strategies, case definitions, or study populations.

This study serves to confirm the importance of several key risk factors for OA in dogs, including age, body weight, and neutering. Chronological age itself is not, of course, modifiable, but the potential to delay the onset of age-associated health conditions, including OA, has been demonstrated in dogs and numerous other species.(27,28) The most successful strategy for achieving this in research animals, severe caloric restriction, is not practical in companion dogs. Such a strategy requires precisely formulated diets and extensive monitoring to prevent malnutrition, and it can actually reduce lifespan in some animals due to nutrition/gene interactions.(48,49) Feeding is also a critical element of the human-animal bond, and asking dog owners to drastically underfeed their canine companions is unlikely to be accepted by many. However, caloric restriction research has done much to elucidate the general mechanisms of aging, raising the possibility that pragmatic therapies to delay aging-associated diseases may be developed.

Body weight is a function of both breed and body condition. Breed risks for specific health conditions, such as OA, can potentially be modified in populations by selective breeding and in individuals by nutritional interventions and, theoretically, the use of drugs targeting the physiologic characteristics of specific breeds that increase disease risk.

Body condition can clearly be altered by nutrition and exercise. Despite the unexpectedly small impact of BCS score on OA risk in this study, OA incidence was greater in overweight and obese dogs than in dogs scored as normal weight. Weight gain after maturity, which likely reflects accumulation of excess adipose, was also a significant risk factor for OA. These findings support the role of obesity as a risk factor for OA and the importance of maintaining optimal body condition in dogs through diet and exercise to minimize OA risk. These interventions are particularly important in larger dogs, who are at greatest risk of obesity and most significantly impacted by weight gain and other risk factors, such as neutering.

Neutering is an important modifiable risk factor for OA. Overall, gonadectomy appears to increase the risk of OA, particularly in larger dogs. Earlier neutering adds to this effect in both large and small dogs. The pathogenesis of OA, of course, is complex and incompletely elucidated, and the role of neutering should be understood in this context, as one of many risk interacting risk factors. Given this complexity, it is not likely that the decision whether or not to neuter an individual dog will be the primary determinant of whether or not that individual develops OA. However, recognizing that an association exists on a population level between neutering and OA risk can help clinicians identify individuals who may be at elevated risk and utilize more proactive surveillance to ensure early diagnosis and appropriate preventative and therapeutic interventions.

Any potential increase in OA risk associated with neutering must also be considered in the context of other risks and benefits associated with neutering on both an individual and population level. Changes in neutering practices intended to mitigate one health risk may entail unpredictable increases in other risks not yet identified, so judicious integration of such information with the larger context and the circumstances and needs of individual patients is critical.

Future research in such large, primary-care cohorts can further clarify the relationship between putative risk factors and common aging-associated diseases such as OA. This will support efforts to reduce the burden of these conditions on companion dogs and their caregivers by informing appropriate preventative intervention.

## Limitations

As a retrospective cohort study, this analysis relied on electronic medical records, collected across thousands of clinics and veterinarians. While these data provide longitudinal insights into risk factors associated with incident diseases, the data themselves were not collected to test hypotheses surrounding risk factors for incident osteoarthritis. The study population (all dogs seen between 1998-2019 and dying in 2019) was large and expected to be representative of the patient population at Banfield during this period, but it may not be representative of the overall U.S. companion dog population.

Electronic health records rely on real-time recording of interactions between clinicians, clients, and patients. Information provided by clients, such as age, breed, and prior medical history, may be subject to misclassification or recall bias. Information provided by veterinarians, such as diagnosis of OA and scoring of BCS, may be subject to misclassification bias and may vary over time and between clinicians. For example, a diagnosis of OA could be based on clinical impression, history and physical examination, radiographic interpretations, or other factors, and no standardized diagnostic criteria were in place.

Overall incidence rates may differ from true population-level incidence, and should be interpreted with caution. Additionally, percentiles were used to summarize adult body weight and BCS across the follow-up window, which fails to capture duration of time over or underweight. Finally, age at desexing was only known for dogs being desexed at a Banfield Pet Hospital clinic, which does not account for desexing occurring outside of Banfield clinics.

## Author Contributions

JG conducted the data analyses and wrote the Materials and Methods and the Results sections. BM wrote the Introduction and Discussion sections. All authors contributed to manuscript revisions and have read and approved the final manuscript.

## Acknowledgements

The authors would like to thank Alexander Naka for significant contributions to the initial conception and design of this project, Zane Koch for assistance with data processing and analysis, Banfield Pet Hospital for sharing the data set analyzed in this project, and the veterinarians and clients of Banfield Pet Hospital for generating this invaluable information resource.

## Contribution to the Field

Osteoarthritis, also called degenerative joint disease, is the most common joint disorder in dogs. It is an incurable, progressive condition that can cause significant pain and disability. Identifying risk factors for the development of osteoarthritis can help veterinarians recognize individual dogs at high risk and make appropriate efforts to diagnose and treat the condition as early as possible. Identifying those risk factors that can be changed can help dog owners and veterinarians prevent or mitigate the development of this condition in some dogs. This study analyzed a very large body of data about companion dogs examined by veterinarians in many general practice veterinary clinics throughout the United States. This analysis confirmed the importance of several previously suspected risk factors for the development of osteoarthritis, including age and body weight. The study also added to existing information about the relationship between the risk of developing osteoarthritis and several factors, including body weight, desexing, and age at desexing. This research will be useful both to clinicians treating canine patients and to other researchers studying the factors that lead to the development of osteoarthritis in dogs.

