## Supplemental figures and tables for "Body weight, gonadectomy, and other risk factors for diagnosis of osteoarthritis in companion dogs"

**Supplemental Materials**

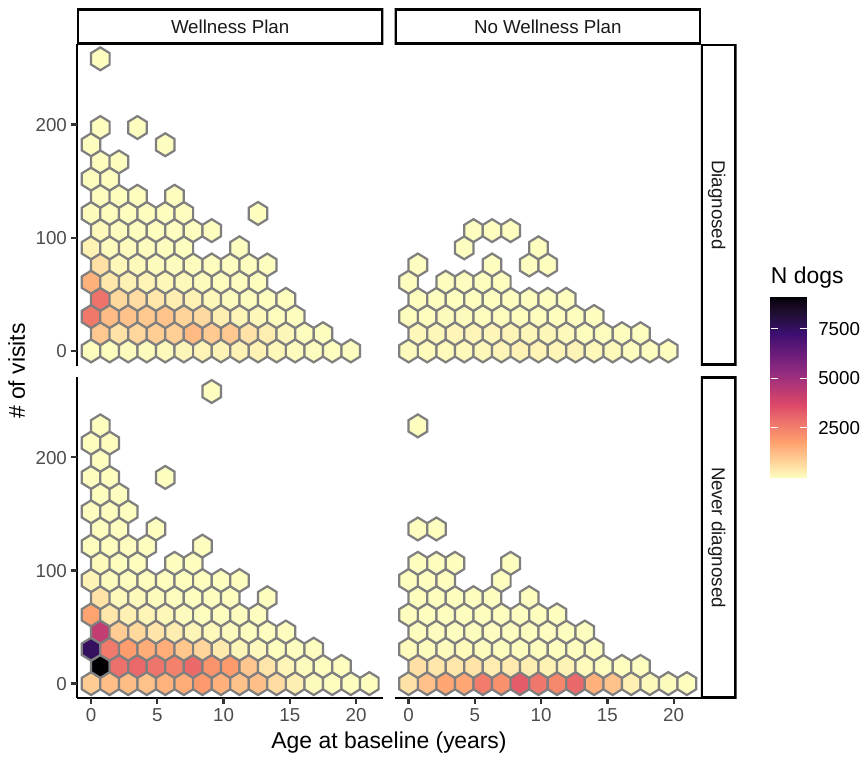

**Supplemental Figure 1.** Hexagonal heatmap of the number of visits reported stratified by the presence or absence of being on a Wellness plan and by OA diagnosis. Color mapping indicates the number of dogs within each bin, where darker values indicate higher numbers of dogs. Dogs on wellness plan have a higher number of visits as well as disproportionally more diagnoses, providing evidence of significant detection bias.

**Supplemental Table 1.** Top three most frequent dog breeds sorted by weight group (cohort quantiles of maximum weight) with sample sizes, OA incidence, and average weight.

| **Breed** | **Total Dogs** | **OA Cases** | **Mean weight (lbs)** | **Weight group** |
| --- | --- | --- | --- | --- |
| Chihuahua | 10,147 | 1,255 (12.37%) | 10.07 | [2,14.6] |
| Yorkshire Terrier | 6,570 | 868 (13.21%) | 9.54 | [2,14.6] |
| Maltese | 3,806 | 554 (14.56%) | 11.09 | [2,14.6] |
| Shih Tzu | 6,251 | 927 (14.83%) | 14.99 | (14.6,29.6] |
| Pug | 2,901 | 689 (23.75%) | 24.56 | (14.6,29.6] |
| Dachshund | 4,355 | 671 (15.41%) | 16.66 | (14.6,29.6] |
| Beagle | 3,560 | 982 (27.58%) | 35.26 | (29.6,66.5] |
| Pit Bull | 5,027 | 935 (18.6%) | 65.63 | (29.6,66.5] |
| Siberian Husky | 1,791 | 590 (32.94%) | 61.55 | (29.6,66.5] |
| Labrador Retriever | 11,718 | 4,721 (40.29%) | 74.22 | (66.5,262] |
| German Shepherd | 4,665 | 1,622 (34.77%) | 76.38 | (66.5,262] |
| Golden Retriever | 3,762 | 1,364 (36.26%) | 75.99 | (66.5,262] |

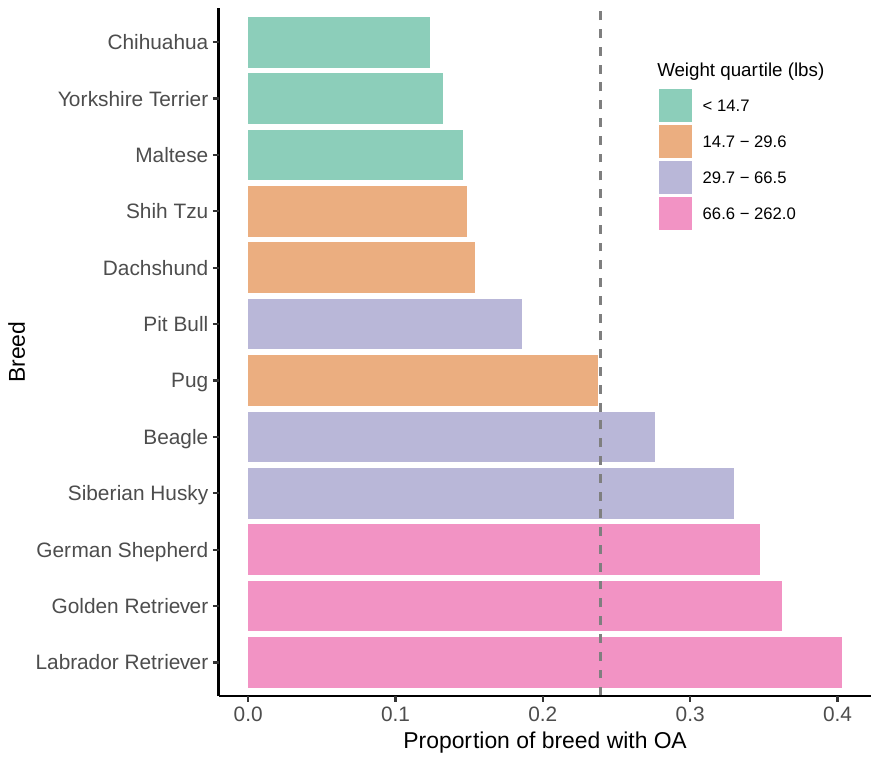

**Supplemental Figure 2.** Barplot showing breed-specific incidence of OA among the 12 representative dog breeds selected for breed-specific sensitivity analysis. Colors reflect weight group assignments, based on the cohort quartiles of maximum body weight. Vertical dashed line is the cohort-specific incidence of OA (23.9%)

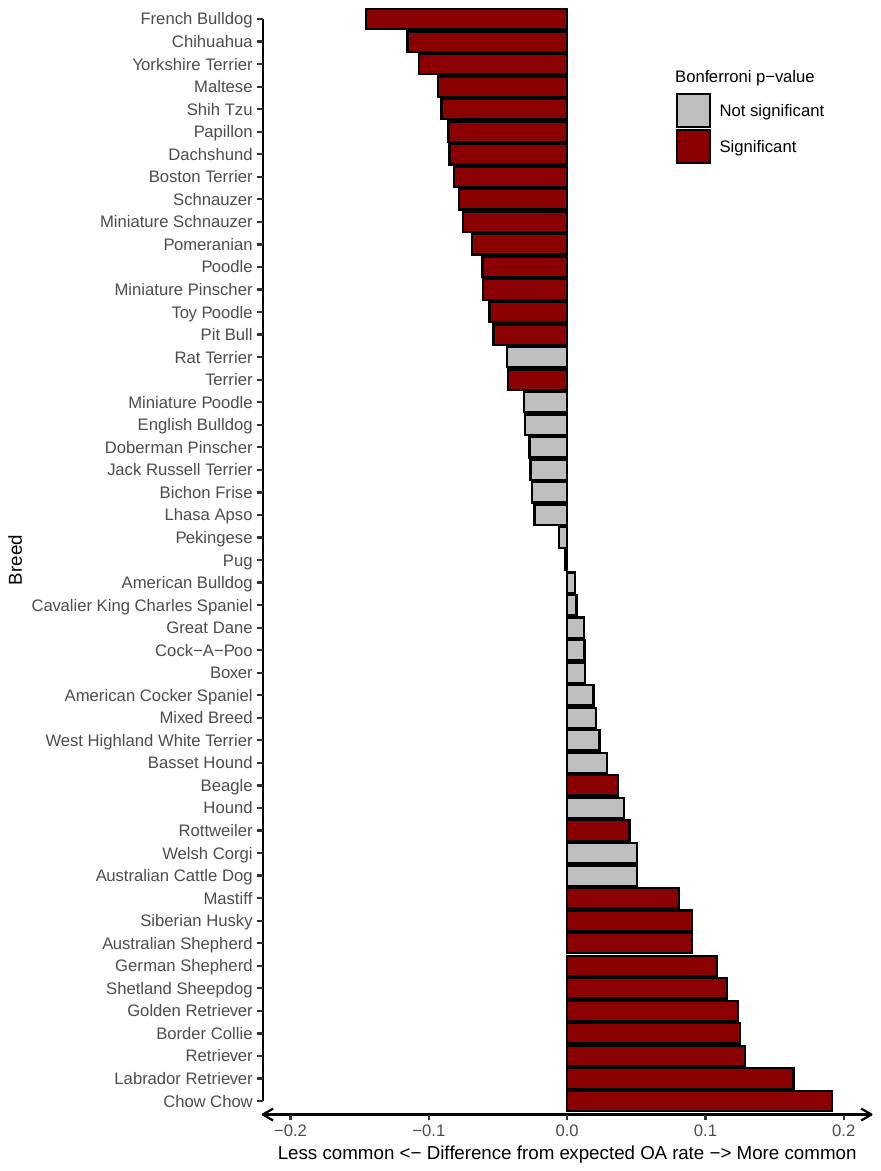

**Supplemental Figure 3.** Barplot showing difference between breed-specific incidence of OA against the cohort-wide incidence of OA for breeds with sample size of at least 500. Color reflects binomial test of proportion is statistically significant, using Bonferroni adjustment.

**Supplemental Table 2.** Breed-specific stratified Cox proportional hazard model testing the associations of primary risk factors on OA-free survival.

| **Breed** | **Variable** | **HR (95% CI)** | **Std. HR (95% CI)** | **p-value** | **Concordance** | **N obs** |
| --- | --- | --- | --- | --- | --- | --- |
| Chihuahua | Age at baseline | 1.31 (1.29-1.33) | 3.40 (3.14-3.68) | <0.001*** | 0.77 | 9,536 |
|  | Max weight | 1.03 (1.02-1.04) | 1.17 (1.11-1.23) | <0.001*** |  |  |
|  | Max BCS | 1.08 (1.02-1.13) | 1.10 (1.03-1.17) | 0.007** |  |  |
|  | Desexed | 1.30 (1.09-1.56) | 1.30 (1.09-1.56) | 0.004** |  |  |
|  | Male | 1.01 (0.90-1.12) | 1.01 (0.90-1.12) | 0.913 |  |  |
| Yorkshire Terrier | Age at baseline | 1.34 (1.31-1.37) | 3.50 (3.19-3.84) | <0.001*** | 0.78 | 6,219 |
|  | Max weight | 1.03 (1.01-1.04) | 1.12 (1.05-1.21) | 0.001** |  |  |
|  | Max BCS | 1.23 (1.14-1.32) | 1.25 (1.16-1.35) | <0.001*** |  |  |
|  | Desexed | 1.09 (0.88-1.34) | 1.09 (0.88-1.34) | 0.431 |  |  |
|  | Male | 1.00 (0.87-1.15) | 1.00 (0.87-1.15) | 0.981 |  |  |
| Maltese | Age at baseline | 1.34 (1.30-1.38) | 3.58 (3.17-4.04) | <0.001*** | 0.77 | 3,601 |
|  | Max weight | 1.02 (1.01-1.03) | 1.08 (1.03-1.15) | 0.004** |  |  |
|  | Max BCS | 1.15 (1.05-1.26) | 1.16 (1.06-1.28) | 0.002** |  |  |
|  | Desexed | 1.14 (0.87-1.50) | 1.14 (0.87-1.50) | 0.347 |  |  |
|  | Male | 1.12 (0.94-1.33) | 1.12 (0.94-1.33) | 0.193 |  |  |
| Shih Tzu | Age at baseline | 1.35 (1.32-1.38) | 3.96 (3.59-4.38) | <0.001*** | 0.78 | 5,920 |
|  | Max weight | 1.03 (1.01-1.04) | 1.13 (1.05-1.21) | 0.002** |  |  |
|  | Max BCS | 1.05 (0.98-1.13) | 1.05 (0.97-1.14) | 0.201 |  |  |
|  | Desexed | 1.04 (0.85-1.27) | 1.04 (0.85-1.27) | 0.711 |  |  |
|  | Male | 1.02 (0.89-1.17) | 1.02 (0.89-1.17) | 0.802 |  |  |
| Pug | Age at baseline | 1.37 (1.33-1.40) | 3.74 (3.34-4.18) | <0.001*** | 0.77 | 2,712 |
|  | Max weight | 1.01 (1.00-1.02) | 1.08 (0.99-1.18) | 0.081 |  |  |
|  | Max BCS | 1.13 (1.03-1.24) | 1.14 (1.03-1.26) | 0.010* |  |  |
|  | Desexed | 0.91 (0.68-1.22) | 0.91 (0.68-1.22) | 0.531 |  |  |
|  | Male | 0.89 (0.76-1.04) | 0.89 (0.76-1.04) | 0.146 |  |  |
| Dachshund | Age at baseline | 1.33 (1.30-1.36) | 3.64 (3.26-4.06) | <0.001*** | 0.78 | 4,095 |
|  | Max weight | 1.02 (1.01-1.04) | 1.18 (1.08-1.29) | <0.001*** |  |  |
|  | Max BCS | 0.98 (0.91-1.06) | 0.98 (0.89-1.07) | 0.649 |  |  |
|  | Desexed | 1.13 (0.85-1.51) | 1.13 (0.85-1.51) | 0.396 |  |  |
|  | Male | 0.93 (0.80-1.09) | 0.93 (0.80-1.09) | 0.361 |  |  |
| Beagle | Age at baseline | 1.34 (1.31-1.37) | 3.48 (3.17-3.81) | <0.001*** | 0.77 | 3,311 |
|  | Max weight | 1.02 (1.01-1.02) | 1.23 (1.15-1.32) | <0.001*** |  |  |
|  | Max BCS | 1.06 (0.99-1.13) | 1.07 (0.99-1.15) | 0.097 |  |  |
|  | Desexed | 1.22 (0.92-1.62) | 1.22 (0.92-1.62) | 0.17 |  |  |
|  | Male | 0.93 (0.82-1.06) | 0.93 (0.82-1.06) | 0.291 |  |  |
| Pit Bull | Age at baseline | 1.37 (1.34-1.40) | 3.39 (3.12-3.69) | <0.001*** | 0.78 | 4,614 |
|  | Max weight | 1.01 (1.00-1.01) | 1.15 (1.07-1.25) | <0.001*** |  |  |
|  | Max BCS | 1.01 (0.95-1.08) | 1.01 (0.93-1.10) | 0.778 |  |  |
|  | Desexed | 1.76 (1.44-2.15) | 1.76 (1.44-2.15) | <0.001*** |  |  |
|  | Male | 1.04 (0.90-1.20) | 1.04 (0.90-1.20) | 0.621 |  |  |
| Siberian Husky | Age at baseline | 1.42 (1.38-1.47) | 4.30 (3.80-4.86) | <0.001*** | 0.81 | 1,641 |
|  | Max weight | 1.01 (1.01-1.02) | 1.27 (1.14-1.41) | <0.001*** |  |  |
|  | Max BCS | 0.98 (0.90-1.07) | 0.98 (0.88-1.09) | 0.672 |  |  |
|  | Desexed | 1.58 (1.10-2.28) | 1.58 (1.10-2.28) | 0.014* |  |  |
|  | Male | 1.10 (0.93-1.31) | 1.10 (0.93-1.31) | 0.262 |  |  |
| Labrador Retriever | Age at baseline | 1.39 (1.38-1.40) | 3.84 (3.68-4.00) | <0.001*** | 0.78 | 10,591 |
|  | Max weight | 1.01 (1.01-1.01) | 1.22 (1.17-1.26) | <0.001*** |  |  |
|  | Max BCS | 1.02 (0.99-1.05) | 1.03 (0.99-1.06) | 0.146 |  |  |
|  | Desexed | 1.29 (1.14-1.46) | 1.29 (1.14-1.46) | <0.001*** |  |  |
|  | Male | 0.94 (0.89-1.00) | 0.94 (0.89-1.00) | 0.067 |  |  |
| German Shepherd | Age at baseline | 1.36 (1.34-1.38) | 3.39 (3.17-3.62) | <0.001*** | 0.77 | 4,258 |
|  | Max weight | 1.02 (1.02-1.02) | 1.44 (1.36-1.52) | <0.001*** |  |  |
|  | Max BCS | 0.93 (0.88-0.97) | 0.91 (0.86-0.97) | 0.002** |  |  |
|  | Desexed | 1.45 (1.21-1.72) | 1.45 (1.21-1.72) | <0.001*** |  |  |
|  | Male | 0.87 (0.79-0.97) | 0.87 (0.79-0.97) | 0.011* |  |  |
| Golden Retriever | Age at baseline | 1.39 (1.36-1.41) | 3.76 (3.48-4.05) | <0.001*** | 0.77 | 3,433 |
|  | Max weight | 1.01 (1.00-1.01) | 1.10 (1.02-1.17) | 0.007** |  |  |
|  | Max BCS | 1.04 (0.98-1.10) | 1.04 (0.98-1.12) | 0.208 |  |  |
|  | Desexed | 1.27 (1.03-1.57) | 1.27 (1.03-1.57) | 0.027* |  |  |
|  | Male | 1.05 (0.93-1.17) | 1.05 (0.93-1.17) | 0.449 |  |  |

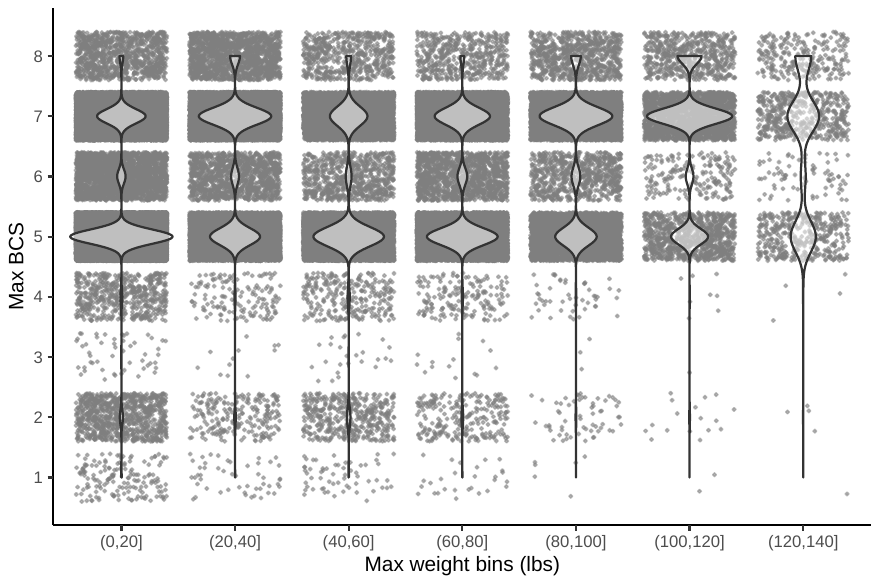

**Supplemental Figure 4.** Grouped scatter and violin plot of the distribution of max BCS scores binned in 20 lb max weight intervals.
